# A Tissue-Bioengineering Strategy for Modeling Rare Human Kidney Diseases *In Vivo*

**DOI:** 10.1101/2021.10.20.465120

**Authors:** J.O.R. Hernandez, X. Wang, M. Vazquez-Segoviano, M.F. Sobral-Reyes, A. Moran-Horowich, M. Sundberg, M. Lopez-Marfil, D.O. Lopez-Cantu, C.K. Probst, G.U Ruiz-Esparza, K. Giannikou, E.P. Henske, D.J. Kwiatkowski, M. Sahin, D.R. Lemos

**Affiliations:** Renal Division, Brigham and Women’s Hospital, Boston, MA 02115; Cancer Genetics Lab, Division of Pulmonary and Critical Care Medicine, Brigham and Women’s Hospital, Boston, MA, 02115; Center for LAM Research and Clinical Care, Division of Pulmonary and Critical Care Medicine, Brigham and Women’s Hospital, Boston, MA, 02115; Rosamund Zander Stone Translational Neuroscience Center, Department of Neurology, Boston Children’s Hospital, MA, 02115; Harvard Medical School, Boston, MA 02115; Harvard Stem Cell Institute, Cambridge, MA 02138

**Keywords:** hiPSCs, organoids, Tuberous Sclerosis Complex, Angiomyolipoma

## Abstract

The lack of animal models for certain human diseases precludes our understanding of disease mechanisms and our ability to test new therapies *in vivo*. Here we generated kidney organoids from Tuberous Sclerosis Complex (TSC) patient-derived-hiPSCs to recapitulate a rare kidney tumor called angiomylipoma (AML). Organoids derived from *TSC2*^*-/-*^ hiPSCs but not from isogenic *TSC2*^*+/-*^ or *TSC2*^*+/+*^ hiPSCs shared a common transcriptional signature and a myomelanocytic cell phenotype with kidney AMLs, and developed epithelial cysts, replicating two major TSC-associated kidney lesions driven by genetic mechanisms that cannot be robustly and consistently recapitulated with transgenic mice. Transplantation of multiple *TSC2*^*-/-*^ kidney organoids into the kidneys of immunodeficient rats allowed us to recapitulate AML and cystic kidney disease *in vivo*, in a scalable fashion and with fidelity, and to test the efficiency of rapamycin-loaded nanoparticles as a novel approach to ablate AMLs by inducing apoptosis triggered by mTOR-inhibition. Collectively, these methods represent a novel tissue-bioengineering strategy for rare disease modeling *in vivo*.

## Introduction

A number of human diseases are driven by genetic and cellular mechanisms that cannot be recapitulated using transgenic mouse models. The lack of an *in vivo* model represents a limitation for mechanistic studies and for testing new therapies on a preclinical level. One such disease is renal angiomyolipoma (AML), a tumor found in 80% of patients with Tuberous Sclerosis Complex (TSC). AML growth can cause kidney failure and lead to premature death due to the formation of vascular aneurysms prone to spontaneous bleeding (1, 2). Rapamaycin analogs (rapalogs) are the main treatment for AMLs. Rapalogs reduce tumor size by inhibiting the dysregulated activity of mTORC1 that results from biallelic inactivation of either *TSC1* or *TSC2* and loss of activity of the encoded proteins, namely TSC1 (hamartin) or TSC2 (tuberin), which repress the metabolic activator kinase mTOR (3). Because rapalogs are mainly delivered orally, these drugs can cause undesired side effects including proteinuria, oral ulcers, dyslipidemia and pneumonitis (4, 5). The lack of animal models is a major obstacle for the development of novel therapies and for the improvement of rapalog-based treatments.

The etiological mechanisms driving AMLs are difficult to recapitulate with transgenic animal models. AMLs are driven by a Knudson’s two-hit mechanism of tumorigenesis that involves a second copy neutral loss-of-heterozygosity (LOH) mutation in the *TSC1* or, more commonly, in the *TSC2* locus, in addition to a pre-existing germline inactivating mutation (6, 7). The second hit results in biallelic inactivation of either *TSC1* or *TSC2*, and constitutive mTOR activation, driving anabolic cell metabolism and aberrant tissue growth (8). Efforts to recapitulate these mechanisms *in vivo* have been challenging due to the fact that biallelic inactivation of *TSC1* or *TSC2* during development causes embryonic lethality. In addition, the cell type giving raise to AML lesions has remained elusive (9, 10), contributing to the lack of success in previous attempts to ablate *TSC1* or *TSC2* by means of tissue-specific Cre-mediated recombination (11, 12). Limited success was achieved using an *in vivo* approach that involved low-dose doxycycline-induced DNA recombination events to ablate *Tsc1* stochastically in mice carrying a ubiquitously expressed Cre transgenic allele (13). Using this strategy, small kidney lesions with characteristics of AMLs could be detected, but the resulting number of mice carrying lesions was small, reducing the suitability of this model for either mechanistic or drug-testing studies (13).

Here we present a novel approach for modeling rare human kidney diseases *in vivo*, using transplanted hiPSC-derived AML kidney organoids. We show that nephric differentiation of patient-derived and genetically-edited *TSC2*^*-/-*^ hiPSCs but not of isogenic *TSC2*^*+/-*^ hiPSCs (14), results in formation of two-dimensional (2D) and three-dimensional (3D) kidney tissues recapitulating TSC-associated AML and cystic disease. Transplantation of multiple 3-D *TSC2*^*-/-*^ organoids into the kidneys of immunodeficient rats resulted in fully vascularized human AML xenografts that allowed us to test the efficacy of an *in situ* approach to rapidly ablate AMLs. The methodology presented here is broadly applicable for the study of other rare kidney diseases for which no *in vivo* experimental model currently exists.

## Results

### *TSC2* inactivation does not impair nephric differentiation of hiPSCs

To understand how lack of TSC2 affects the early stages of nephric differentiation, we used an *in vitro* protocol previously developed for the directed differentiation of hiPSCs into kidney tissues (15). We used a set of three isogenic hiPSC lines that included a line derived from an individual carrying a heterozygous 9-bp deletion in the *TSC2* locus (*TSC2*^+/−^), a second isogenic *TSC2*^-/-^ hiPSC line that was generated by introducing a TALEN-engineered second deletion in the wild type (WT) *TSC2* allele of the patient-derived hiPSC line (14), and lastly a *TSC2*^+/+^ hiPSC line in which the original *TSC2* deletion present in the patient-derived hiPSC line was corrected using CRISPR-Cas9 (Figure 1a) (14). The *TSC2*^-/-^ hiPSC line displayed significantly increased mTOR activity compared to the *TSC2*^+/−^ and *TSC2*^+/+^ hiPSC lines, as indicated by the levels of phosphorylation of the ribosomal protein S6 (Phospho-S6) detected by Western Blot (Figure 1b). No differences in phospho-S6 levels were observed between the *TSC2*^+/−^ and *TSC2*^+/+^ hiPSC lines, a finding that was consistent with the idea that one functional *TSC2* allele is sufficient to preserve mTOR activity under these conditions (Figure 1b).

**Figure 1.**
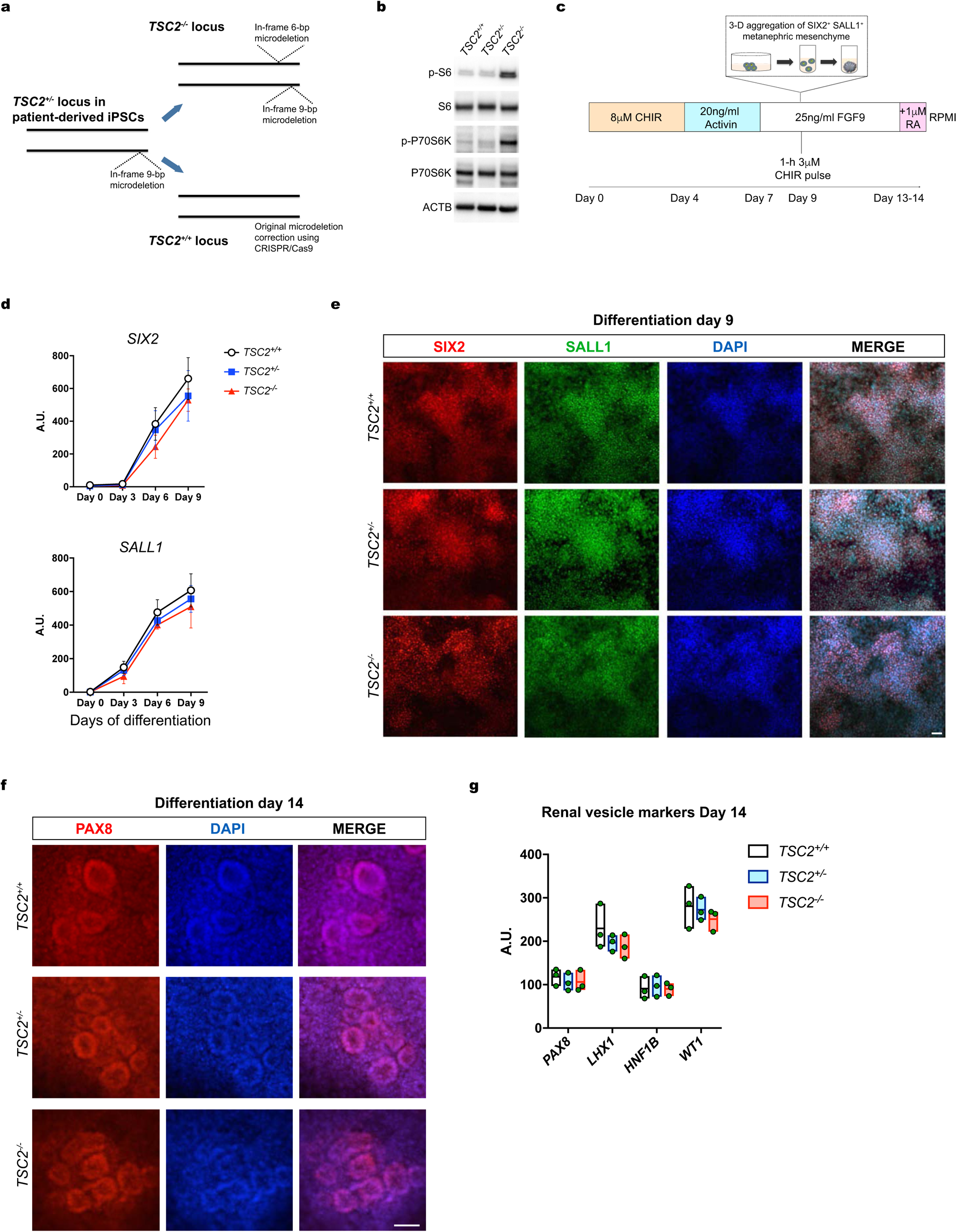
Loss of TSC2 does not impair hiPSC nephric differentiation. **a** Schematic representation of the gene-editing strategies used to either introduce a second inactivating mutation in the wild type allele of patient-derived *TSC2*^*+/-*^ iPSCs, or to correct the original deletion mutation. **b** Immunoblots showing phosphorylation of S6 and p-70S6K in *TSC2*^*+/+*^, *TSC2*^*-/-*^ and *TSC2*^*+/-*^ hiPSCs, indicating mTORC1 activation in the *TSC2*^*-/-*^ cells. **c** Schematic representation for the hiPSC nephric differentiation protocol used to generate 2-D renal tissues and 3-D renal organoids. **d** Quantitative RT-PCR analysis of *SIX2* and *SALL1* mRNAs during the phase of metanephric mesenchyme induction. Curve points represent mean ± SD in arbitrary units (A.U. normalized to β-actin), n = 3 independent technical replicates. **e, f** Representative immunofluorescence images showing SIX2 and SALL1 (**e**), and PAX8 (**f**) in 2-D *TSC2*^*+/+*^, *TSC2*^*-/-*^ and *TSC2*^*+/-*^ cell cultures on Day 9 (**e**) and Day 14 (**f**) of differentiation. **g** Quantitative RT-PCR analysis of *PAX8, LHX1, HNF1B* and *WT1* mRNAs in 2-D *TSC2*^*+/+*^, *TSC2*^*-/-*^ and *TSC2*^*+/-*^ cell cultures on differentiation Day 14, showing similar expression independent of genotype. Floating bars graph represent mean ± SD, n = 3 independent experiments. Scale bars, 50μm.

To generate renal tissues from the three hiPSC lines in 2-D conditions, we used a previously established protocol that involves an initial step of metanephric mesenchyme induction by initial incubation with the WNT activator CHIR99201 (CHIR) at 8μM for four days, followed by incubation with 20ng/ml Activin A for three days (Figure 1c) and subsequent incubation with 25ng/ml fibroblast growth factor 9 (FGF9) for two days (15). On Day 9, the 2D cultures received a 1-h pulse of 3μM CHIR. The 8-μM concentration of CHIR during the initial induction phase was chosen based on comparable mRNA expression profiles observed for the nephric lineage homeobox genes *SIX2* and *SALL1* in the three hiPSC genotypes between Day 0 and Day 9, which were determined by quantitative polymerase chain reaction (qPCR) (Figure 1d). Both SIX2 and SALL1 were also detected by immunofluorescence in cultures of the three hiPSC lines, confirming induction of these nephric lineage markers (Figure 1d). Microscopic field quantification indicated that ∼82%, 85% and 91% of *TSC2*^+/-^, *TSC2*^-/-^ and *TSC2*^+/+^ cells, respectively, were SIX2^+^; and ∼80%, 83% and 87% of *TSC2*^+/-^, *TSC2*^-/-^ and *TSC2*^+/+^ cells, respectively, were SALL1^+^, indicating efficient induction of all three cell lines (Figure 1e). After confirming induction of metanephric mesenchyme from the three hiPSC lines, we assessed their ability to subsequently form the renal vesicles that give rise to nephrons. To that end, we continued incubating the metanephric mesenchyme cultures with FGF9 until Day 14. During this stage, the cultures were exposed to a 24-h pulse of 1μM retinoic acid on Day 13 of differentiation (Figure 1c). This modification to the original protocol was adopted from a recent protocol for the differentiation of podocytes (16) and was introduced based on our observation of glomeruli of an increased size present in Day 21 cell cultures (Supplementary figure 1). In order to assess the formation of renal vesicles on Day 14 we used a combination of immunostaining for the homeobox transcription factor PAX8 and qPCRs to detect the expression of the nephric homeobox genes *PAX8, LHX1, HNF1* and *WT1*. We confirmed the formation of cell aggregates with the morphology of renal vesicles containing PAX8^+^ renal progenitor cells (Figure 1f). In addition, we confirmed similar expression levels for renal vesicle markers in *TSC2*^-/-^, *TSC2*^+/-^ and *TSC2*^+/+^ cell cultures, and found that all were upregulated compared to undifferentiated hiPSC cultures (Figure 1g).

Taken together, these results indicated that dysregulated mTOR activity is not an impediment for the nephric specification of hiPSCs into metanephric mesenchyme and the formation of renal vesicles under 2-D culture conditions.

### *TSC2*^*-/-*^ hiPSCs generate 3-D renal organoids with characteristics of kidney AMLs *in vitro*

After confirming the successful formation of renal vesicles in 2-D cultures for all three hiPSC lines, we proceeded to induce the formation of 3-D kidney organoids. Following the original published procedure (17) we dissociated the metanephric mesenchyme cell cultures on Day 9, to obtain single cell suspensions that were subsequently aggregated on ultra-low attachment tissue culture surfaces. The subsequent differentiation process was the same as the one used in our 2-D cultures (Figure 1c). This procedure resulted in the formation of organoids that we harvested on Day 21 and interrogated for the presence of several known kidney AML markers, the most common renal abnormality observed in TSC patients (18, 19). We first investigated the expression and distribution of premelanoma 17 (PMEL) using the mouse monoclonal antibody HMB45, which is clinically used for pathologic diagnosis (20, 21). Our immunofluorescence analysis did not detect PMEL^+^ cells in *TSC2*^+/-^ and *TSC2*^+/+^ kidney organoids (Figure 2d), a finding that was consistent with a previous study using the Morizane protocol for nephric differentiation of hPSCs (22). By contrast, we identified a large population of PMEL^+^ cells in *TSC2*^-/-^ organoids, covering an area that was approximately 6.2% of the organoid sections analyzed (Figure 2a). PMEL^+^ cells were present in the interstitial space, but were not peritubular, a characteristic that distinguished them from renal fibroblasts observed in both human kidneys and hPSC-derived organoids. PMEL^+^ cells had a spindle-like morphology with a granular staining pattern that resembled the characteristic phenotype of human kidney AML cells (Figure 2a, Supplementary Figure 2a) (23). Co-expression of PMEL and glycoprotein NMB (GPNMB), another AML cell marker (24), further confirmed the melanocytic identity of *TSC2*^-/-^ organoid cells (Figure 2b). Next, we investigated the expression and distribution of cells expressing actin alpha 2, smooth muscle (ACTA2, commonly known as alpha smooth muscle actin, another major AML marker (23) that is expressed at low levels in non-injured kidneys and WT organoids (25). We observed that ACTA2 was highly expressed in *TSC2*^-/-^ organoids, with ACTA2^+^ cells covering an area that represented 21.7% of the sections analyzed, compared to 3.28% and 3.7% of the areas in *TSC2*^+/+^ and *TSC2*^+/-^ organoid sections, respectively (Figure 2c). Similar to what we observed for PMEL^+^ cells, ACTA2^+^ cells were sparsely distributed throughout the stroma of *TSC2*^-/-^ organoids (Figure 2c). Consistent with what has been previously reported, the ACTA2^+^ cells observed in *TSC2*^+/-^ and *TSC2*^+/+^ organoids were mostly confined to regions surrounding nephron epithelial structures (Supplementary figure 2b). Morphologically, ACTA2^+^ cells found in *TSC2*^-/-^ organoids had a rounder and myoid morphology that best matched the well-characterized morphology of ACTA2^+^ cells observed in kidney AMLs (Figure 2c, Supplementary figure 2a) (10). By contrast, ACTA2^+^ cells in *TSC2*^+/-^ and *TSC2*^+/+^ organoids presented an elongated spindle-like cell body with extended body processes resembling the morphology of kidney fibroblasts (Supplementary Figure 2b) (25). Given that co-expression of melanocyte and mesenchymal cell markers is unique to AML cells, we next investigated whether *TSC2*^-/-^ organoid cells had a myomelanocytic phenotype. Using immunofluorescence we were able to establish that ∼94% of GPNMB-expressing cells were ACTA2^+^ (Figure 2d). Cells co-expressing both markers were also detected in AML samples, but not in activated fibroblasts/myofibroblasts from injured *TSC2*^*+/+*^ organoids. In order to validate the myomelanocytic cell phenotype using a different technical approach, we analyzed the distribution of PMEL^+^ and ACTA2^+^ cell populations in *TSC2*^-/-^ organoids by means of fluorescence activated cell sorting (FACs). Our analysis identified a population of ACTA2^+^ cells that represented approximately 31.2% of the total cells recorded in *TSC2*^-/-^ organoid samples (Figure 2e, Supplementary figure 4). The same cell population constituted 3.5% and 4.1% of the cells recorded in *TSC2*^+/-^ and *TSC2*^+/+^ organoid samples, respectively (Figure 2e). Within the ACTA2^+^ cell population, approximately 24.3% of the cells also expressed PMEL (Figure 2e), whereas no PMEL^+^ ACTA2^-^ cells were detected, a finding that was consistent with our immunofluorescence results for GPNMB and ACTA2. Taken together, our histology and FACS data identified a population of *TSC2*^-/-^ organoid cells that recapitulates the unique phenotype observed in some populations of AML tumor cells. Lastly, we confirmed the abnormal expression of GPNMB and ACTA2 in *TSC2*^-/-^ organoids by means of Western Blot performed on protein extracts from *TSC2*^+/-^ and *TSC2*^+/+^ and *TSC2*^-/-^ organoids (Figure 2f).

**Figure 2.**
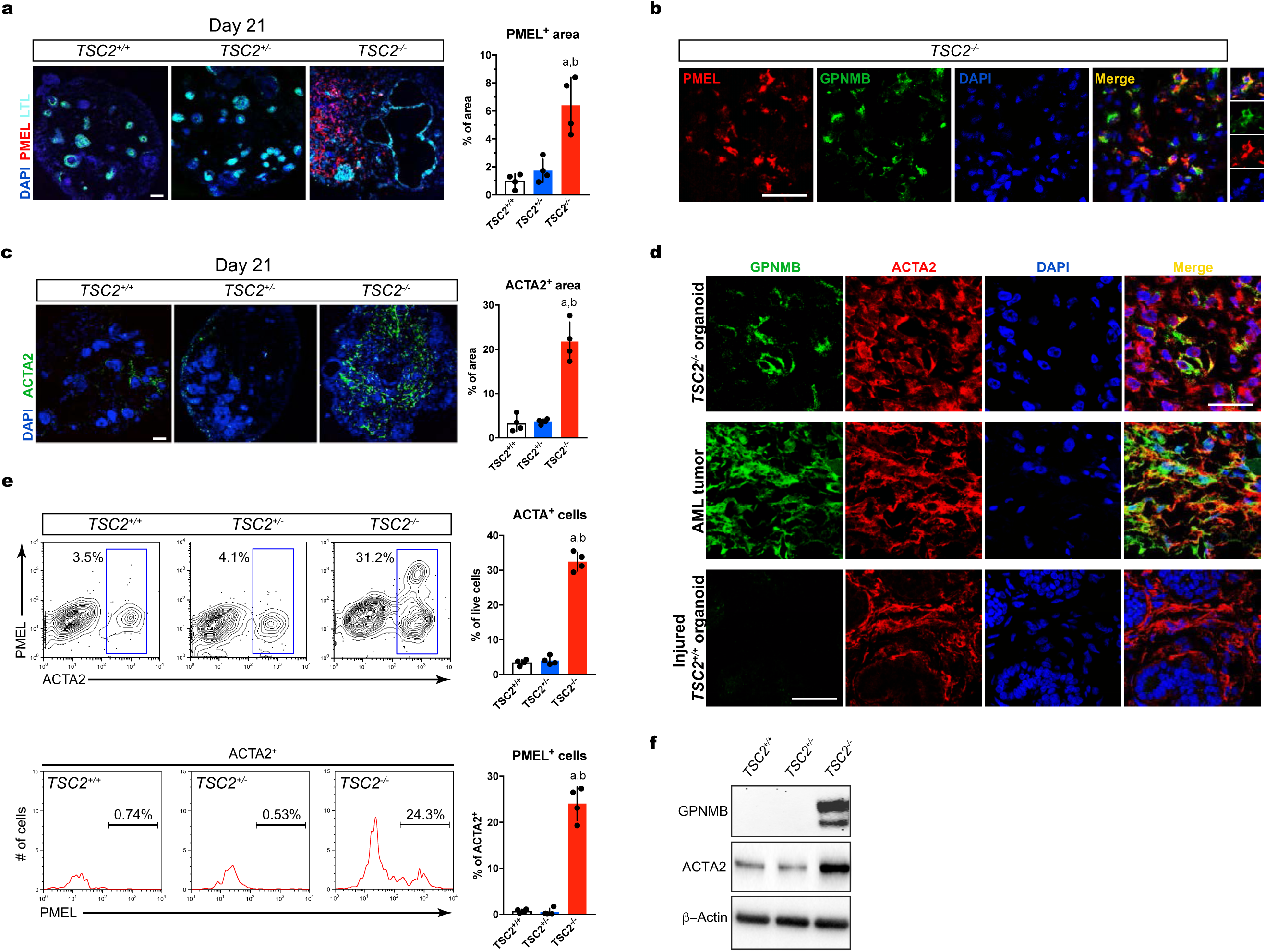
*TSC2*^*-/-*^ renal organoids recapitulate human AML tumor phenotype. **a, b, c** Confocal optical sections showing LTL with PMEL (**a**), PMEL and GPNMB (**b**), and ACTA2 (**c**) in Day 21 3-D renal organoids derived from *TSC2*^*+/+*^, *TSC2*^*-/-*^ and *TSC2*^*+/-*^ hiPSCs. Bar graphs in (**a**) and (**b**) represent mean ± SD, n = 4 experiments. a, *P*<0.05 *TSC2*^*-/-*^ *vs. TSC2*^*+/+*^, b, *P*<0.05 *TSC2*^*-/-*^ *vs. TSC2*^*+/-*^. **d** Representative confocal immunofluorescence sections showing expression of GPNMB and ACTA2 in Day-21 *TSC2*^*-/-*^ renal organoids, kidney AML tumor and fibrotic Day-28 *TSC*^*+/+*^ renal organoids injured by incubation with interleukin 1β for 96h. **e** Representative FACs plots and quantification showing the percentage of ACTA2^+^ cells and PMEL^+^ cells in Day-21 *TSC2*^*+/+*^, *TSC2*^*-/-*^ and *TSC2*^*+/-*^ renal organoids. Bar graphs represent mean ± SD, n = 4 experiments. a, *P*<0.01 *TSC2*^*-/-*^ *vs. TSC2*^*+/+*^, b, *P*<0.01 *TSC2*^*-/-*^ *vs. TSC2*^*+/-*^. **f** Representative immunoblot showing expression of GPNMB and ACTA2 in renal organoids from the three genotypes. Scale bars 50μm.

Collectively these *in vitro* data indicate that renal organoids derived from *TSC2*^-/-^ hiPSCs, but not from *TSC2*^+/-^ hiPSCs, possessed several key characteristics of renal AML tumors, including the presence of myomelanocytic cells with typical AML morphology. These results also provide critical experimental support to the hypothesis that AML cells derive from renal progenitor cells during development.

### Gene expression analysis of *TSC2*^*-/-*^ organoids validates an AML identity

In order to investigate how loss of *TSC2* affects gene expression in hiPSC-derived renal organoids, we performed whole transcriptome RNA sequencing (RNA-seq) of *TSC2*^+/-^, *TSC2*^+/+^ and *TSC2*^-/-^ organoids. We used the normalized read counts (FPKM) to perform pairwise DESeq2 differential gene expression analyses, to compare *TSC2*^-/-^ organoids to *TSC2*^+/+^ and *TSC2*^+/-^ organoids (www.qlucore.com). Several hundred genes were upregulated or downregulated at a false discovery rate (FDR)/ q-value < 0.05, p-value < 0.002, |log_2_ fold| > 2 and |log_2_ fold change| <-2 (Supplementary Table 1). Among the genes upregulated in the *TSC2*^-/-^ organoids were: *MLANA, PMEL, GPNMB, MITF, CTSK* and *ACTA2*, with median expression fold change 121x-, 88x-, 34x-, 21x-, 14x-, 9.5x, respectively, all of which are known to be highly expressed in the renal angiomyolipoma (Figure 3a). In contrast, DEseq2 analysis of *TSC2*^+/+^ *vs. TSC2*^+/-^ renal organoids did not reveal remarkable differences. Multiple housekeeping genes, including *GPDH, GUSB, RPL19*, showed no significant difference or minimal changes in expression across the 3 kidney organoid genotypes providing further confirmation that the changes observed were specific to the genes identified (Supplementary figure 3a).

**Figure 3.**
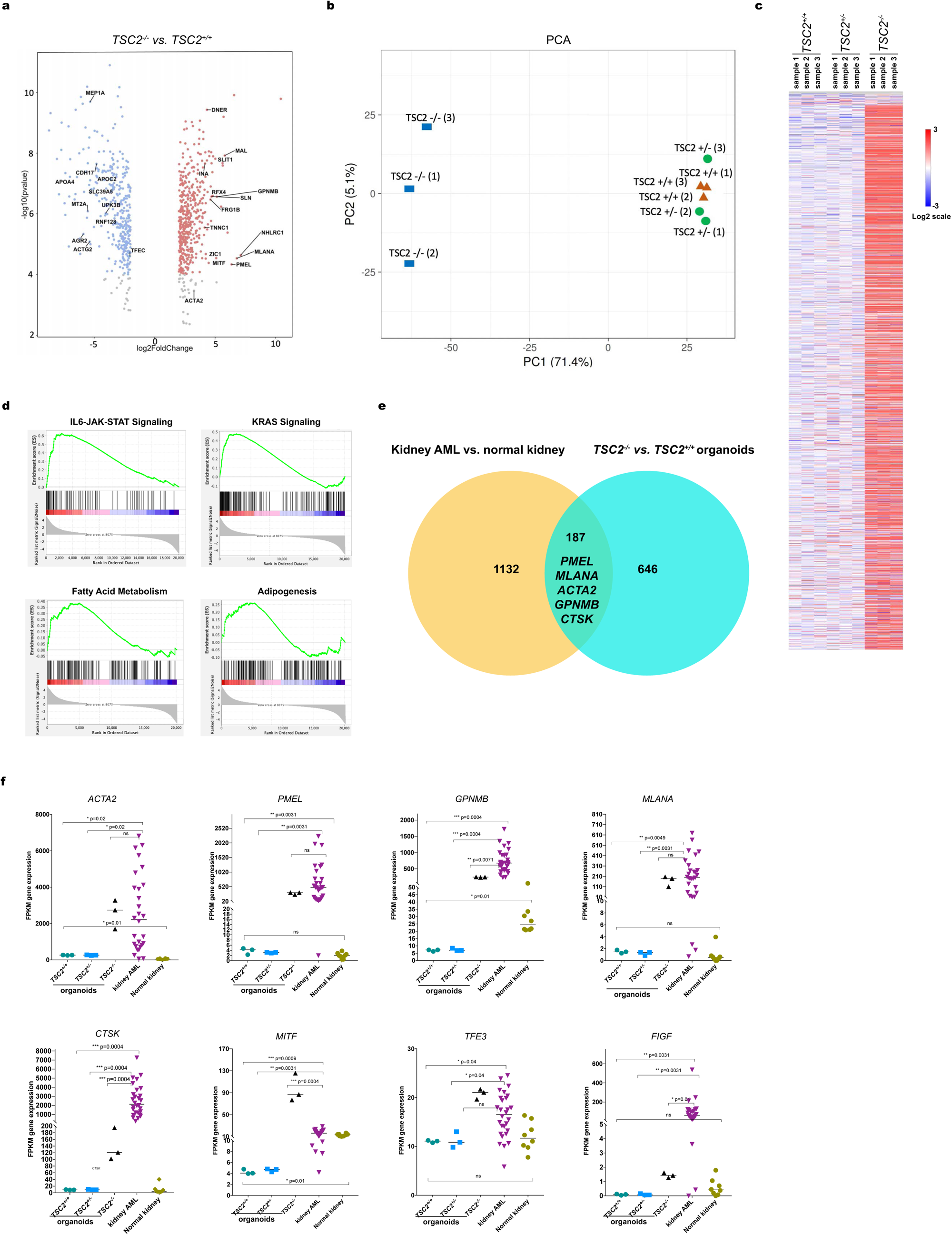
Gene expression analysis of *TSC2*^*-/-*^, *TSC2*^*+/-*^, and *TSC2*^*+/+*^ renal organoids. **a** Volcano plots showing the distribution of all differentially expressed genes (DEGs) in *TSC2*^*-/-*^ renal organoids compared to *TSC2*^*+/+*^ (left) and *TSC2*^*+/-*^ (right) renal organoids (FDR<0.05). Each dot represents a unique gene; red denotes log_2_ (fold change) > 2, up-regulated genes in *TSC2*^*-/-*^; blue denotes log_2_ (fold change) <-2, down-regulated in *TSC2*^*-/-*^. Selected statistically significant upregulated and downregulated genes (NCBI/Entrez names) are indicated. **b** Principal Component Analysis of RNA-Seq data from renal organoids of the three genotypes, n=3 samples for each genotype, 5 organoids *per* sample. **c** Heatmap showing hierarchical clustering of three different genotypes of kidney organoids using the top 3000 most variable genes. **d** Representative enrichment plots corresponding to GSEA for pairwise comparison of *TSC2* ^*-/-*^ *vs. TSC2*^*+/-*^. **e** Venn diagrams indicating 187 common differentially expressed genes, including signature AML markers, in *TSC2*^*-/-*^ *vs. TSC2*^*+/+*^ renal organoids and kidney AML *vs*. normal kidney. **f** Box plots for a panel of AML marker genes for *TSC2*^*-/-*^, *TSC2*^*+/+*^ and *TSC2*^*+/-*^ renal organoids (n=3 each) compared to human kidney AML (n=28) and human kidney (n=8). P-values are selected comparisons are shown. Gene expression is shown in FPKM values.

Principal Component Analysis (PCA) of RNA-Seq data for all three genotypes also demonstrated that the *TSC2*^*-/-*^ organoids were transcriptionally different from *TSC2*^+/+^ and *TSC2*^+/-^ kidney organoids, which clustered together (Figure 3b). Hierarchical clustering using Spearman rank correlation for the 3,000 most variable genes also demonstrated that the *TSC2*^*-/-*^ organoids were distinct from the other two genotypes, with the latter appearing similar (Figure 3c). Gene set enrichment analysis (GSEA) was performed comparing *TSC2*^*-/-*^ renal organoid RNA data to that of *TSC2*^*+/+*^ renal organoids, and showed enrichment in multiple gene sets at FDR/q<0.25, with similar results in comparison of *TSC2*^*-/-*^ and *TSC2*^*+/-*^ renal organoids. Hallmark gene sets enriched in expression in *TSC2*^*-/-*^ renal organoids in each comparison included IL6-JAK-STAT3 signaling, adipogenesis, angiogenesis, fatty acid metabolism, KRAS signaling and estrogen response (Figure 3d, Supplementary figure 3a, b).

We then investigated the similarity between the genes identified in the comparisons between *TSC2*^*-/-*^ *vs. TSC2*^*+/+*^organoids against differential genes identified in the comparison between kidney AML *vs*. normal kidney (Figure 3e). Remarkably, the 187 common differentially expressed genes included five of the six genes listed above, as well as *TFE3* and *FIGF*. Notably, *MITF* is not among these shared genes, as its expression was not significantly increased in AMLs in comparison to normal kidney, although its transcriptional activity is markedly increased and accounts for the increased expression of *MLANA, PMEL, GPNMB*, and *CTSK* (26).

Overall, our gene expression analysis indicated that nephric differentiation of *TSC2*^*-/-*^ iPSCs leads to marked expression differences compared to similar differentiaton of *TSC2*^*+/+*^ and *TSC2*^*+/-*^ iPSCs, and that many of the genes whose expression is increased are also upregulated in renal AML, confirming that this differentiation protocol leads to formation of an *TSC2*^*-/-*^ AML-like organoid.

### *TSC2* inactivation drives renal organoid tubule cyst formation

Based on the dilated appearance of putative proximal tubules in *TSC2*^*-/-*^ 3-D organoids (Figure 2a), we set out to study the phenotype of *TSC2*^*-/-*^ nephrons. We noted that brightfield imaging of Day-21 2-D *TSC2*^*+/+*^ and *TSC2*^*+/-*^ cell cultures contained tubule like structures, whereas cultures derived from *TSC2*^-/-^ hiPSCs showed cavitated structures resembling cysts (Figure 4a). Because cysts are another common abnormality found in TSC patients with renal manifestations (27), we investigated the cellular composition of the cysts to determine whether they were formed by tubule epithelial cells (TECs) derived from *TSC2*^*-/-*^ hiPSCs. By means of immunofluorescence, we detected nephron segments corresponding to distal tubules expressing cadherin 1 (CDH1), proximal tubules containing brush borders labeled by lotus tetranoglobus lectin (LTL) and glomeruli containing podocytes expressing podocalyxin 1 (PODXL) in the cell aggregates derived from *TSC2*^*+/+*^ and *TSC2*^*+/-*^ hiPSCs (Figure 4b). In both cases, the segments looked anatomically normal and were sequentially connected, therefore indicating continuous nephrons (Figure 4b). In the case of *TSC2*^*+/-*^ cell aggregates, the cystic structures stained positive for both CDH1 and LTL, indicating the cyst-like structures could comprise distal tubule and/or proximal tubule cells (Figure 4c). Cyst formation occurred with a frequency of ∼4.7 cysts/well for *TSC2*^-/-^ cultures, compared with 0 and ∼0.5 cysts/well for *TSC2*^+/+^ and *TSC2*^+/+^ cultures, respectively (Figure 4b). Of note, 2-D cyst-like structures developed despite the fact that *PKD1* expression without addition of cAMP modulators such as forskolin (Figure 4h), a method previously employed for the induction of cysts in kidney organoid models of polycystic kidney disease (PKD) (28). Another observable phenotype in 2-D TSC2-/-nephrons was that PODXL^+^ cells presented a less compact appearance, and were loosely grouped, suggesting dysmorphic glomerular structures (Figure 4c).

**Figure 4.**
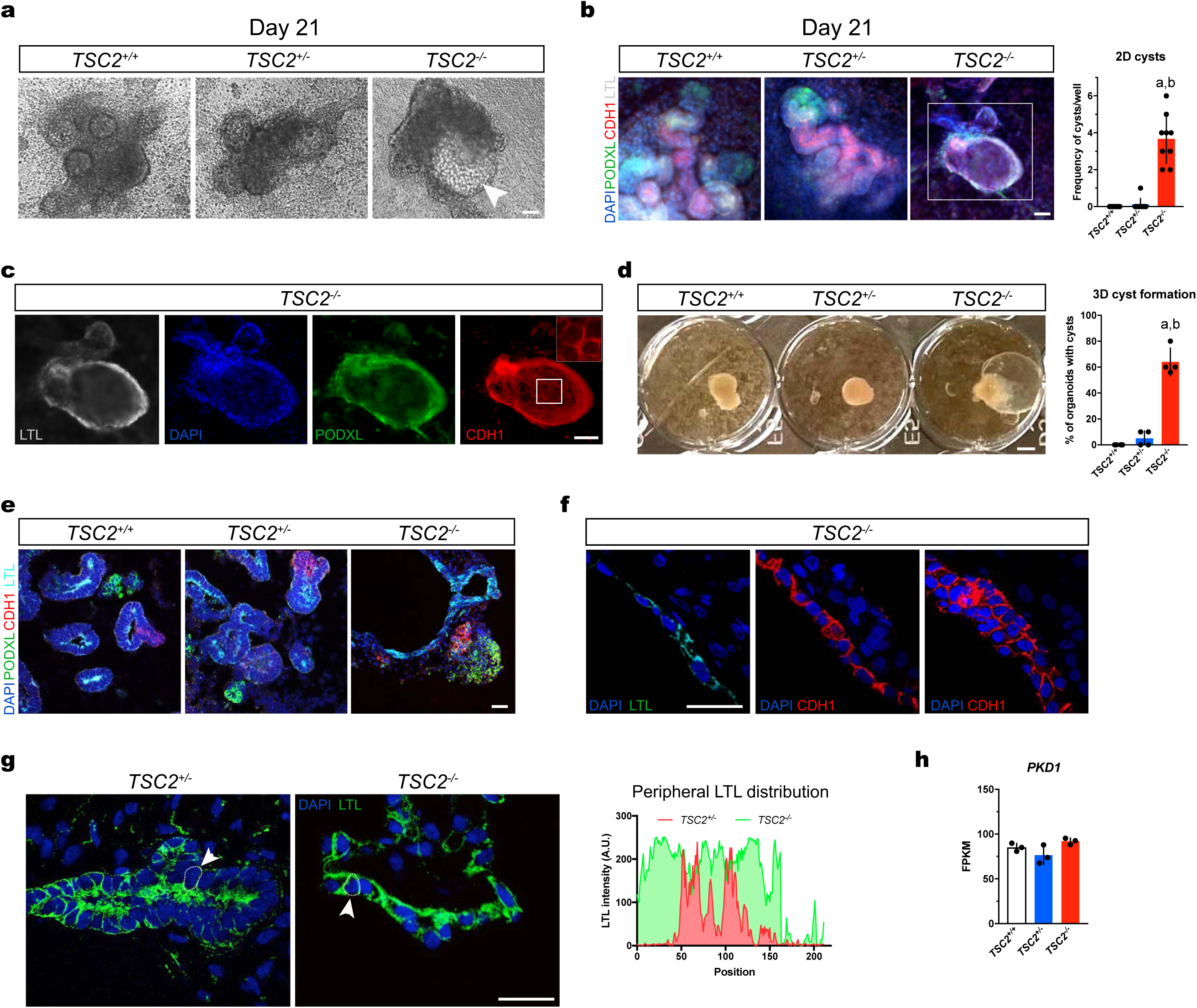
*TSC2* inactivation drives cystogenesis during nephric differentiation. **a** Representative brightfield images of 2-D *TSC2*^*-/-*^ hiPSC-derived renal tissues on Day 21 of differentiation. The arrowhead indicates a visible cyst. **b** Immunofluorescence image showing 2-D nephrons derived from the three hiPSC genotypes, labeled with PODXL1 (glomerulus), LTL (proximal tubule), and CDH1 (distal tubule). Number of cysts per well was quantified. Bar graphs represent mean ± SD, n = 8 wells from 2 independent experiments. **c**, The cyst framed in the *TSC2*^*-/-*^ nephron micrograph in (**a**) is stained for nephron markers by immunofluorescence with inset showing the distal tubule epithelium. **d** Brightfield image of 3-D organoids of the three genotypes showing cystic *TSC2*^*-/-*^ organoids, with quantification of cyst formation. Bar graphs represent mean ± SD, n = 3 experiments. \**P*<0.05; \*\**P*<0.01. **e** Representative confocal sections showing organoid glomeruli, proximal and distal tubule regions as indicated by PODXL1, LTL and CDH1 staining. A cyst associated with a proximal tubule is shown in the *TSC2*^*-/-*^ organoid section. **f** Anatomical organization of *TSC2*^*-/-*^ organoid cyst lining visualized using confocal imaging, both PTEC and DTEC single-cell layers and DTEC multicellular regions are shown with LTL and CDH1 immunofluorescence. **g** Representative confocal imaging showing the alterations in polarity observed in PTECs lining *TSC2*^*-/-*^ organoid cysts, compared to *TSC2*^*+/-*^ organoid proximal tubule, using LTL staining. A representative quantification of LTL signal distribution in the periphery of individual cells is shown. Scale bars, 50μm (**a**,**b**,**c**,**e**), 25μm (**f, g**) and 1mm (**d**).

These observations prompted us to analyze the spatial the analysis of spatial arrangements of cystic nephron structures in 3-D organoids. On Day 18 of the 3D differentiation protocol, we observed protruding dome-like structures indicating incipient cyst formation on the surface of *TSC2*^-/-^, but not of *TSC2*^+/+^ or *TSC2*^+/-^ differentiating spheroids by Day 16 (Supplementary fig 5a). On Day 21, individual cysts growing out of the *TSC2*^-/-^ AML organoids were clearly observed by brightfield microscopy (Figure 4d). Similar to what has been reported in organoid models of polycystic kidney disease, we found the size of cysts to be larger in 3-D aggregates grown on low-attachment surfaces compared to 2-D cell cultures (Supplementary figure 5a). Nephron segment analysis using immunofluorescence microscopy confirmed that *TSC2*^+/+^ and *TSC2*^+/-^ metanephric mesenchyme aggregates had formed kidney organoids as indicated by the detection of properly segmented nephrons in which CDH1^+^ distal tubules were connected with LTL^+^ proximal tubules, and PODXL^+^ nephrons, on Day 21 of the differentiation protocol (Figure 4e). In *TSC2*^-/-^ AML organoids, the analysis confirmed that the cells conforming the lining of the cysts stained positive for CDH1 and/or LTL, but not for PODXL, indicating that similar to what is observed in TSC patients (29) the cysts are specifically associated with tubular nephron segments, but can also emerge from multiple segments (Figure 4f, Supplementary figure 5b). The immunofluorescence analysis also showed that while mainly individual tubule segments were involved, cysts lined by a combination of both proximal and distal tubule cellular components were also observed in *TSC2*^-/-^ AML organoid cysts, possibly representing transition regions (Figure 4f). Unlike the single-cell layers observed lining the cyst regions associated with well-defined tubule segments, the mixed-cell lining regions were multi-layered (Figure 4f), resembling the columnar epithelium that can be observed in the renal cysts of TSC patients (29). The distribution of LTL staining, representing polarity of brush borders, revealed another aspect of TSC cystic disease, namely loss of cell polarity in the single-cell proximal tubule stretches of cyst lining, in contrast to the polarized cells observed in *TSC2*^+/-^ tubules (Figure 4g). Of note, cyst formation in *TSC2*^-/-^ AML organoids did not involve loss of polycystin 1 (PKD-1) expression, as indicated by the similar mRNA levels detected in our organoid RNAseq analysis (Figure 4h).

These findings indicated that *TSC2*^*-/-*^ renal organoids recapitulate key features of cystic kidney disease associated with TSC, and support the concept that renal epithelial cyst formation is also driven by loss of *TSC2*.

### Transplanted *TSC2*^-/-^ AML organoids recapitulate TSC-associated kidney AML and cystic disease *in vivo*

After determining that our *in vitro TSC2*^-/-^ hiPSC-derived renal organoids recapitulated major features of AMLs TSC-associated cystic disease, we sought to investigate the effect of vascularization on phenotype development. The paucity of vascularization is a major limitation of *in vitro* hPSC-derived kidney organoid models (30). We transplanted Day-18 *TSC2*^-/-^ pre-organoid stage spheroids into the kidneys of 10-week-old immunodeficient RNU rats (Figure 5a, b). We sought to establish how *in vivo* vascularization alters the anatomy and growth of *TSC2*^-/-^ AML organoids, and whether this approach would result in an animal model capable of recapitulating TSC-associated renal abnormalities. Our strategy took advantage of the size of RNU rat kidneys, which allowed for the simultaneous transplantation of two ∼500-μm Day-18 spheroids (Figure 5a, b). In addition, we transplanted organoids into both kidneys of each rat, giving us the ability to compare xenografts with different genotypes in the same animal.

**Figure 5.**
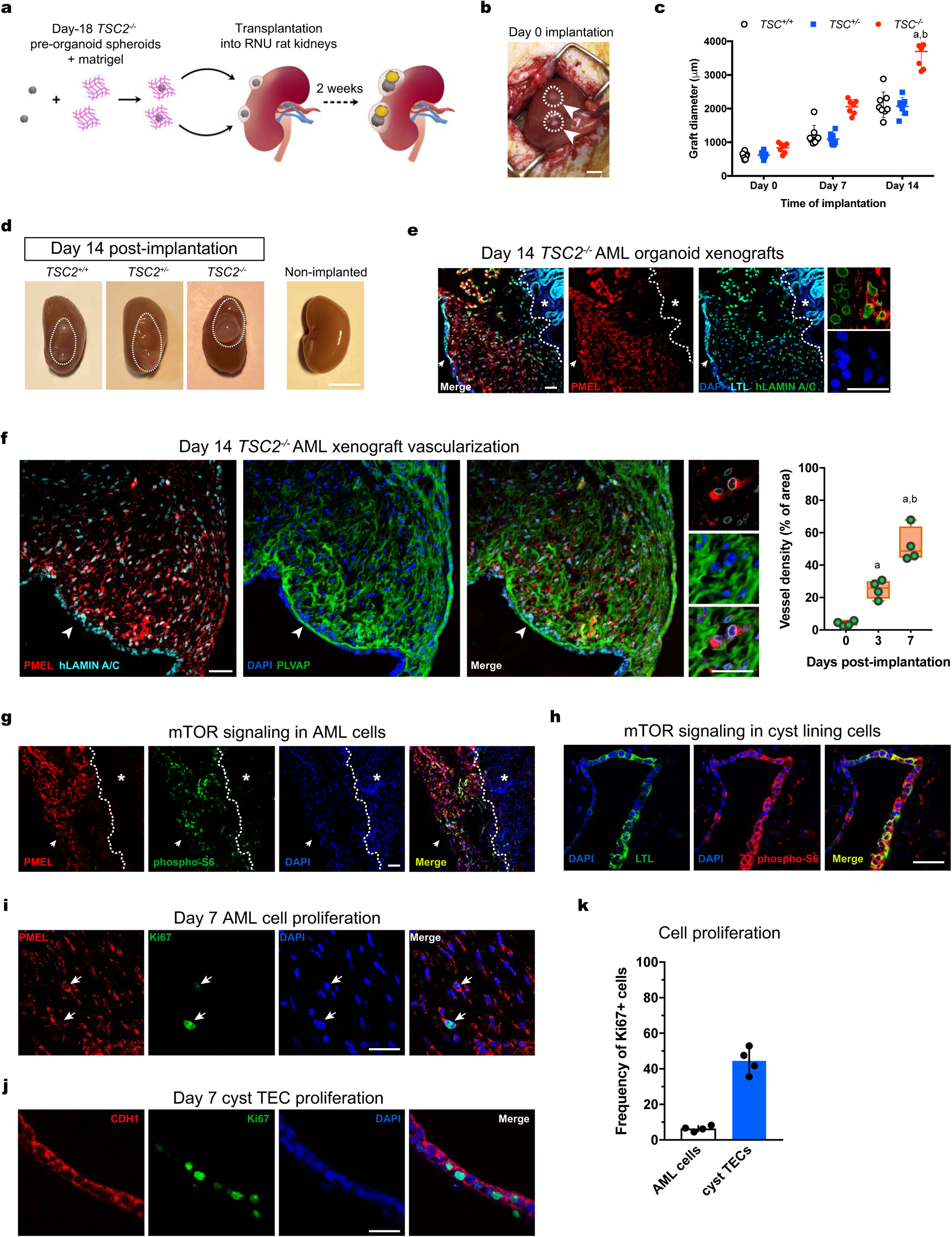
Orthotopic *TSC2*^*-/-*^ renal organoid xenografts model TSC-associated AML and TSC cystic disease *in vivo*. **a** Schematic depicting the strategy used for the transplantation of Day 18 pre-organoid spheroids in the subcapsular region of RNU rat kidneys. **b** Represeantative image showing two spheroids implanted on Day 0, each indicated by an arrowhead. **c** Quantification of graft size over the period of two weeks. Scatter dot plot shows mean ± SD, n = 8 grafts, 4 rat kidneys. \**P*<0.05; \*\**P*<0.01. d Representative image of RNU rat kidneys with engrafted *TSC2*^*+/+*^, *TSC2*^*-/-*^ and *TSC2*^*+/-*^ organoids on Day 14 post-transplantation. **e** Representative confocal sections of *TSC2*^*-/-*^ AML grafts showing wide distribution of human PMEL^+^ cells and LTL^+^ PTECs forming the cyst lining, the latter indicated by a white arrow. LTL-stained rat proximal tubules are indicated by a white star. **f** Representative immunostaining and confocal images showing the thorough vascularization indicated by PLVAP^+^ vessels, observed in close apposition to PMEL^+^ cells. Quantification of vessel density shown in the floating bar graph for grafts harvested on Day 0, Day 3 and Day 7 post-transplantation. Values represent mean ± SD, n = 4 grafts. \**P*<0.05; \*\**P*<0.01. **g, h** Representative confocal immunofluorescence images for phospho-S6 in PMEL^+^ cells (**g**) and cyst-lining PTECs (**h**) in *TSC2*^*-/-*^ AML xenografts. In (**g**) the white arrow indicates the luminal compartment, while the white star indicates the rat kidney. **i, j** Detection of Ki67 in PMEL^+^ cells (**i**) and PTECs (**j**) in *TSC2*^*-/-*^ AML grafts on Day 7 post-transplantation. The white arrows indicate Ki67-expressing human PMEL^+^ cells. Quantification of PMEL^+^ Ki67^+^ and LTL^+^ Ki67^+^ cells shown in the bar graph representing mean ± SD, n = 4 grafts. \**P*<0.05. Scale bars 1cm (**b**), 2cm (**d**), 50μm (**e**,**f**,**g**,**h**), 25μm (**e**,**f** high magnification panels), 25μm (**i**,**j**).

Assessment of xenograft size indicated a significantly higher growth rate for *TSC2*^-/-^ AML organoids (4.43 fold change), compared to *TSC2*^+/-^ (3.55 fold change) or *TSC2*^+/+^ kidney organoids (3.4 fold change) at Day 14 post-transplantation (Figure 5c). At Day 14, cysts could be seen on the surface of kidneys containing *TSC2*^-/-^ AML organoids but not in kidneys carrying the *TSC2*^+/-^ or *TSC2*^+/+^ organoids (Figure 4d). At this time point, we surgically removed the rat kidneys and examined the xenografts. To distinguish human organoid tissues from rat tissues by means of immunofluorescence microscopy, we used antibodies against either the human nuclear antigen (HNA) or the human isoform of Lamin A/C (hLamin A/C). Analysis of AML markers revealed the widespread and abundant presence of human cells expressing either ACTA2 or PMEL in the *TSC2*^-/-^ xenografts, but not in *TSC2*^+/-^ or *TSC2*^+/+^ organoid xenografts, indicating that the AML phenotype was maintained after engraftment (Figure 5e). Both cell types were observed in close proximity to thoroughly distributed microvascular endothelium of host origin that was visualized through detection of the marker plasmalemma vesicle-associated protein (PLVAP) (Figure 5f), which is expressed by the microvasculature of the kidney (31). In addition, mTOR activation, assessed by pS6 expression, was observed in both ACTA2^+^ or PMEL^+^ cells of *TSC2*^-/-^ AML organoid graft cells, but not in the adjacent normal rat tissue or in *TSC2*^+/+^ or *TSC2*^+/-^ organoid xenografts, indicating that metabolic activation was consistent with xenograft growth (Figure 5g). Analysis of the epithelium in *TSC2*^-/-^ xenografts confirmed the formation of cystic tubules that were lined by both PTECs and DTECs in varying proportions, arranged into stretches of single-cell lining, that alternated with stretches presenting a multilayered organization, as observed in *TSC2*^-/-^ organoids *in vitro* (Figure 4f). Similar to the TECs found in the *in vitro* cysts of *TSC2*^-/-^ AML organoids, the TECs present in the in graft cysts lacked polarity and had activated mTORC1 signaling. Similar to the AML cells, activation of mTORC1 activation was also observed in epithelial cyst cells of the *TSC2*^-/-^ grafts (Figure 5h). Despite the common activation of mTORC1 observed in both AML cells and TECs, most proliferative activity occurred in the cysts lining (Figure 5i, j, k), suggesting that graft growth was mainly a consequence of cyst hyperplasia. This result is consistent with both the lack of proliferative activity observed in classic kidney AML cells (32), and the neoplastic activity observed in TSC-associated kidney cyst lining epithelial cells (29, 33).

Collectively our transplantation experiments showed that introduction of a vascularization enhanced the phenotype of *TSC2*^-/-^ AML organoids, permitting for the first time the recapitulation of human TSC-associated AML *in vivo* with a high degree of anatomical and molecular fidelity.

### Delivery of rapamycin-loaded nanoparticles abrogates orthotopic *TSC2*^-/-^ AML xenografts

Our results with transplanted *TSC2*^-/-^ AML organoids in RNU rats prompted us to use this new animal model to examine the potential benefit of a novel formulation for delivery of the mTOR inhibitor Rapamycin, a well-known therapy used for the treatment of AML tumors in TSC patients (27). To this end, we designed an experiment in which Rapa was encapsulated in ultra-fine solid biopolymeric nanoparticles and delivered into the subcapsular space adjacent to the organoid grafts (Supplementary figure 6a-e). We devised this approach to enable targeted delivery of small doses of rapamycin locally over a systemic delivery approach which is known to result in undesired drug effects in other organs. We confirmed the absorption of rapa-loaded nanoparticles by kidney organoids *in vitro*, by visualizing the uptake of nanoparticles loaded with Bodipy 650/665-X (Supplementary figure 6). Incubation of kidney organoids with nanoparticles loaded with 500ng Rapa, *in vitro*, resulted in progressive loss of tissue viability within 48h (Supplementary figure 6f, g). Based on these results, we tested two doses of Rapa for the *in vivo* treatments, namely 500ng and 2μg, which we injected under the kidney capsule approximately 2mm away from each *TSC2*^-/-^ AML organoid xenograft, 2 weeks after transplantation. We repeated the treatment 3 days later to complete a regimen of two injections, and harvested the whole kidneys one week after the first injection (Figure 6a). We measured the size of the organoid xenografts 3 days and 1 week -i.e. at tissue-collection time- after the first injection. Our measurements showed that rapamycin nanoparticle treatment resulted in a dramatic reduction of organoid xenograft size (Figure 6b). Immunofluorescence staining showing Caspase 3 (Casp3) activation in AML ^+^ cells suggested that the treatment induced cell apoptosis (Figure 6c). Western blotting of organoid xenografts showed increased cleaved caspase 3 at 3 and 7 days of treatment (Figure 6e).

**Figure 6.**
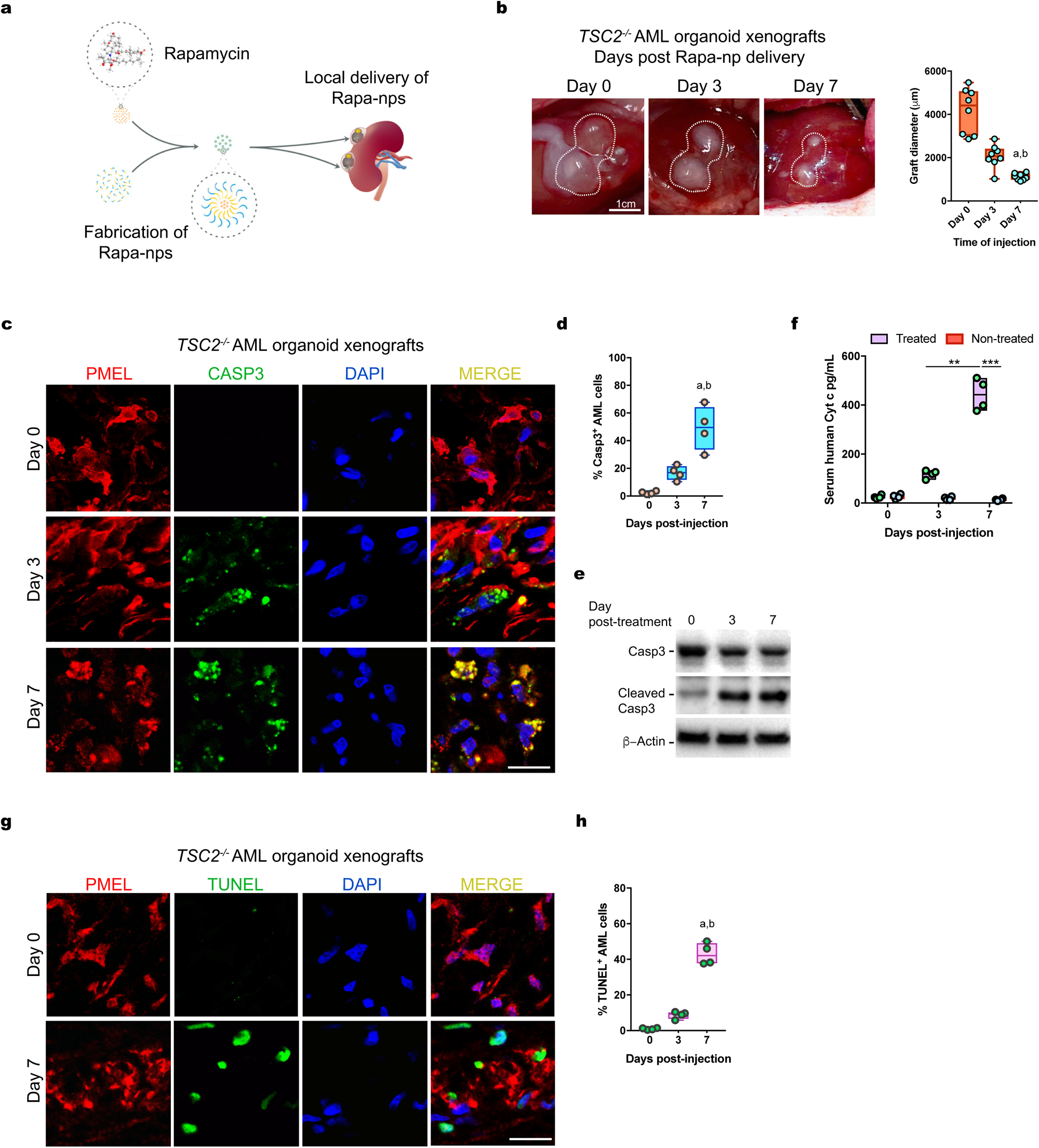
Ablation of *TSC2*^*-/-*^ AML organoid xenografts treated with Rapamycin-loaded nanoparticles. **a** Schematic showing the strategy for the delivery of Rapa-nanoparticles locally, near the *TSC2*^*-/-*^ AML organoid xenografts. **b** Representative photographs showing the size of *TSC2*^*-/-*^ xenografts on Day 3 and Day 7 post Rapa-np delivery. Quantified diameter values on the floating bar graph represent mean ± SD, n = 8 grafts. \**P*<0.05; \*\**P*<0.01. **c** Representative immunofluorescence images for the detection of activated Casp3 in *TSC2*^*-/-*^ AML organoid xenograft PMEL^+^ myoid cells on Day 3 and Day 7 post drug delivery. **d** Floating bar graph with quantification of PMEL^+^ Casp3^+^ cells on Days 3 and 7 post drug delivery. Mean ± SD are reported, n = 4 grafts. \**P*<0.05; \*\**P*<0.01. **e** Detection of Cleaved Casp3 on Day 0 (free organoid), 3 and 7 in protein extracts from *TSC2*^*-/-*^ AML xenografts (Day 3 and 7). **f** Quantification of human Cytochrome C in the serum of RNU carrying rats *TSC2*^*-/-*^ AML organoid xenografts, treated with Rapa-np and controls that did not receive the treatment. Floating bar graph shows mean ± SD, n = 4 rats per group. \**P*<0.05; \*\**P*<0.01.**g** Representative confocal imaging depicting detection of DNA fragmentation by TUNEL on sections of *TSC2*^*-/-*^ AML xenografts, three and seven days post Rapa-np delivery. **h** Floating bar graph for the quantification of PMEL^+^ TUNEL^+^ cells shows mean ± SD are reported, n = 4 grafts per group. \**P*<0.05; \*\**P*<0.01. Scale bars 1cm (**b**), 25mm (**c**,**g**).

**Figure 7.**
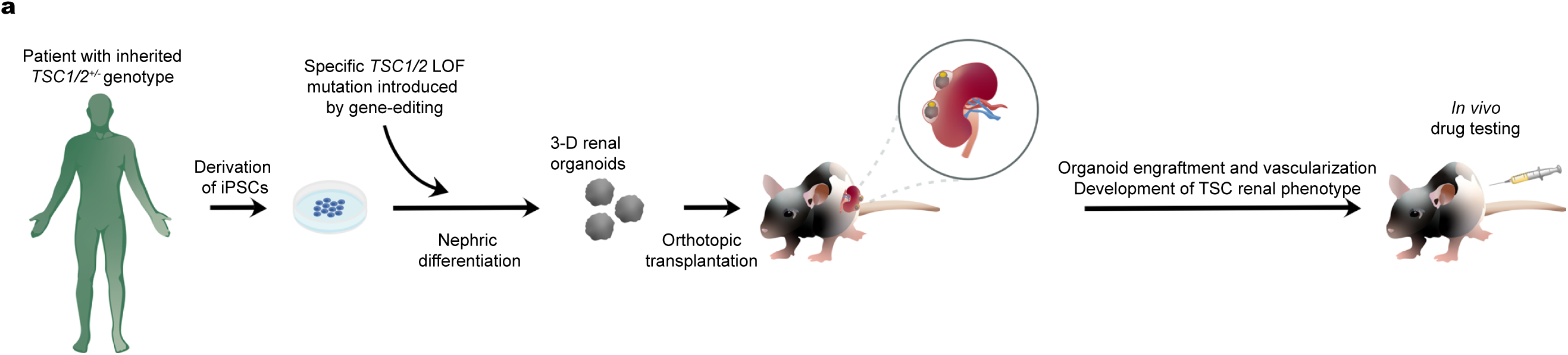
Proposed preclinical animal model for the study of human TSC-associated renal diseases with drug-testing capabilities. **a** Schematic representation of the strategy used in this study depicting the generation of 3-D renal tissues from patient-derived hiPSC to which an LOF mutation were introduced by gene-editing, resulting in complete loss of TSC2. Transplantation of spheroids allows maturation of AML tissues and cystic tubules and vascularization *in vivo*, for the recapitulation of renal manifestations driven by loss of TSC2.

In addition to the analysis of Casp3 activity, we detected increasing DNA fragmentation in PMEL^+^ cells of treated *TSC2*^-/-^ AML organoid xenografts by means histological analysis of terminal deoxynucleotidyl transferase dUTP nick end labeling (TUNEL) (Figure 6e). Both Casp3 activity and DNA fragmentation indicated that delivery of Rapa-nps was an effective treatment for local ablation of *TSC2*^-/-^ AML organoid xenografts.

Taken together, these results highlight the *TSC2*^-/-^ AML organoid xenografts in RNU rats as a powerful preclinical model for testing drugs and new therapeutic approaches for the treatment of AMLs.

## Discussion

Here we report an experimental strategy for the generation of *in vivo* models of human renal diseases associated with rare genetic disorders that cannot be reproduced in all their complexity with current experimental animal models. Using a combination of genome editing and hiPSC techniques we show that nephric differentiation of hiPSCs carrying biallelic LOF mutations in the *TSC2* locus generates renal tissues that reproduce key anatomical and molecular aspects of human AML tumors. To the best of our knowledge this is the first recapitulation of TSC-associated renal diseases using hiPSCs.

The AML identity of PMEL^+^, ACTA2^+^ *TSC2*^*-/-*^ organoid cells is supported by two critical findings, namely the mixed mesenchymal and melanocyte identity, and cell morphology (34, 35). Melanocytes do not express ACTA2, as indicated by single cell transcriptome profiles of skin cells (36, 37). Similarly, mesenchymal cells in normal kidney tissues do not express melanocyte markers. Both our flow cytometry and histology results show that ACTA2^+^ mesenchymal cells present in *TSC2*^*+/-*^ or *TSC2*^*+/+*^ kidney organoids did not express melanocytic markers, a result that is consistent with single cell transcriptome profiles of WT kidney organoid mesenchymal cells (37). Lastly, in terms of morphology, *TSC2*^*-/-*^ organoid AML cells show a distinct plump spindle-like morphology with that is commonly observed in AML tumor cells (21, 34), and which clearly differs from the morphology of kidney fibroblasts characterized by elongated spindle-shaped cell bodies ending in prolonged processes, as well as from the multipolar dendritic morphology of skin melanocytes (38).

*TSC2*^*-/-*^ AML organoids also recapitulated the epithelial cysts observed in TSC patients. Cystic kidney disease is the second most frequent renal manifestation of TSC and can be triggered by the biallelic inactivation of *TSC1/TSC2* alone (33, 39) or as the result of deletion events that also affect the *PKD1* locus located adjacently in chromosome 16 (40). Our RNAseq data showing that levels of *PKD1* mRNA in *TSC2*^*-/-*^ AML organoids were comparable to mRNA levels in *TSC*^*+/-*^ and *TSC*^*+/+*^ kidney organoids, indicated that the cystogenic mechanisms are solely driven by loss of *TSC2*. Cyst formation can also be associated with a subtype of AML called AML containing epithelial cysts (AMLECs) (41). In the case of AMLECs, the mechanisms of cystogenesis are not well understood, and a role for tumor cells, possibly through the secretion of paracrine factors, has been suggested (41). While those mechanisms have not been investigated in this study, the presence of both an AML phenotype and epithelial cysts is congruent with the idea that early developmental LOH events affect a broader spectrum of kidney tissues, therefore resulting in multiple RAs (8, 27).

Critically, our experiments in RNU rats represent a major step toward modeling TSC-associated renal manifestations *in vivo* using transplanted hiPSC-derived kidney organoids. The rat kidney critically provided vascularization and supported the growth of *TSC2*^*-/-*^ organoids, promoting the hyperproliferative activity of cyst lining epithelial cysts, consistent with the clinical notion that factors present in the kidney and the blood play an important role in the development of cystic kidney disease. We have shown that this model can increase the throughput of disease modeling studies by supporting multiple organoid xenografts that can be used for mechanistic interrogation and for drug-testing. The combination of genome editing approaches for patient-derived hiPSCs and *in vivo* disease modeling using organoid xenografts is a promising strategy for personalized medicine.

## Methods

### hiPSC line maintenance

The hiPSC cell lines 77-patient, 77-TSC2-null and control 77-TSC2WT were a kind gift from Dr. Sahin. The 77-patient line carrying a heterozygous 9-bp deletion in the *TSC2* locus was derived from the lymphoblasts of a patient with TSC using the sendai-virus method for the delivery of *OCT3/4, KLF4, L-MYC, SOX2* and *LIN28* into the cells as previously described (14). The 77-TSC2-null line was generated by introducing a second microdeletionof 6-bp in the WT allele of the 77-patient line, using transcription activator-like effector nucleases TALEN (14). The 77-TSC2WT line was created by correcting the heterozygous microdeletion of TSC2 in 77-patient cell line with CRISPR-cas9 method (14). The three hiPSC lines were maintained in mTeSR™1 medium (STEMCELL Technologies # 85850), in 6-well tissue culture plates (Falcon, #353046) coated with 1% vol/vol LDEV-Free hESC-qualified Matrigel (Corning Life Sciences # 354277) in a 37°C incubator with 5% CO2. Cells were passaged at a 1:3 split ratio once a week, using Gentle Cell Dissociation Reagent (GCDR, STEMCELL Technologies #07174). Each cell line was maintained below passage 45 and mycoplasma contamination was not detected. Permission for the use of hiPSC lines was granted to DRL by our institutional review board (IRB) through IBC protocol 2020B000020.

### hiPSC differentiation

hiPSCs maintenance cultures were washed once with PBS (Life Technologies, #10010-049) and dissociated into single cell suspensions using Accutase (STEMCELL Technologies, #07920). One hundred and fifty thousand cells were plated onto 24-well tissue culture plates (TPP, #92024) coated with 1% Matrigel in mTeSR™1 medium supplemented with ROCK inhibitor molecule Y27632 (10μM) (TOCRIS, #1254). Nephric differentiation was performed as described previously (15), briefly after 24 h, the mTeSR™1 medium was replaced by basic differentiation medium consisting of Advanced RPMI 1640 (Life Technologies, #12633-020) containing 1X L-GlutaMAX (Life Technologies, #35050-061) supplemented with 8 μM CHIR99021 (Sigma-Aldrich, #SML1046) for 4 days. Cells were then cultured in Advanced RPMI + 1X L-GlutaMAX + Activin A (10 ng/ml) (R&D, #338-AC-050) for 3 days. For induction of metanephric mesenchyme the media was replaced with Advanced RPMI + 1X L-GlutaMAX + FGF9 (20 ng/ml) (R&D, #273-F9-025/CF) for 7 days, with a 1-h pulse of 3μM CHIR added on day 9. To generate 3-D organoids, the cells were lifted using Accutase on day 9 and the cell suspensions incubated with CHIR pulse medium 37 °C, 5% CO_2_ for 1h then switched to basic differentiation medium + FGF9 and plated at 50,000 cells *per* well onto ultra-low-attachment plates (Corning, #7007). The plates were centrifuged at 1,500 r.p.m. for 15s, and the cells then cultured 37°C, 5% CO_2_ until day 13. At day 13, a pulse of 1μM retinoic acid was added to the medium containing FGF9, and the organoid or cell cultures were incubated for 24h. At day 14, the cultures where transferred to basic differentiation medium with no additional factors for 7–14 days, and harvested at day 21-28.

### Induction of organoid fibrosis with IL-1β

Day-21 *TSC2*^*+/+*^ organoids were incubated in Advanced RPMI + 1X L-GlutaMAX with 50ng/ml human IL1-β (Sigma-Aldrich # H6291) for 96hs, as previously described (25). The medium containing IL1-β was replaced every 48h.

### Quantitative RT-PCR

Total RNA was extracted using the RNeasy Plus MiniKit (Qiagen). Purity was determined byA260–A280. cDNA was synthesized using oligo(dT) and random primers (iScript Reverse Transcription Supermix; Biorad). Quantitative PCR was performed using the QuantStudio 7 Flex Real-Time PCR System (ThermoFisher Scientific) using TaqMan Gene Expression Assays (ThermoFisher Scientific #4331182). The specific primer pairs used in quantitative PCR are listed in Supplemental Table 2.

### Western blot

*In vitro* and grafted organoids were lysed in RIPA buffer containing Halt Protease and Phosphatase Inhibitor Cocktail (Thermo FisherScientific), using a glass Dounce homogenizer (DWK Life Sciences Kimble™) on ice. Following protein quantification, SDS-PAGE gel run and western blotting were performed as described (25). The following antibodies Cell Signaling Technology were used: anti-ribosomal protein S6 (#2217), anti-phospho S6 (#2211), anti-phospho P70S6K (#9205), anti-P70S6K (#9202), anti-β Actin (#4967), anti-GPNMB (#38313). Anti-ACTA2 was from Sigma-Aldrich (#F3777), anti-human cleaved Caspase 3 was purchased from Abcam (ab2302). Primary antibodies were detected with peroxidase-conjugated anti-rabbit or anti-mouse IgG (1:3000) and visualized with SuperSignal West Femto Substrate (ThermoFisher Scientific #34094). Blots were quantifi ed using ImageJ (https://imagej.nih.gov/ij/).

### Immunofluorescence

Organoids and grafts were fixed with 4% paraformaldehyde in PBS for 30 min in a 96-well plate, washed three times in PBS, then incubated with 30% sucrose (w/w) overnight at 4°C. The next day the tissues were mounted into frozen blocks with O.C.T. compound (Fisher Scientific, #23-730-571) and were cut into 12-μm sections onto glass slides (Fisher Scientific #22-037-247). The sections were washed three times for 5 min in PBS, then incubated in blocking buffer containing 0.15% Triton X-100 and 5% normal donkey serum) for 1 h, followed by incubation with primary antibodies in antibody dilution buffer (0.15% Triton X-100 and 1% BSA in PBS) for 2 h or overnight, then washed three times in PBS. Incubation with secondary antibodies in antibody dilution buffer was performed for 1 h, followed by three washes in PBS and coverslips mounted on Vectashield with DAPI (Vector Labs #H-1200). The primary antibodies used were: anti-SIX2 (Proteintech, cat. no. 11562-1-AP), anti-SALL1 (Abcam #ab41974), anti-PAX8 (Proteintech, cat. no. 10336-1-AP), anti-PMEL (Agilent Technologies, #M063429-2), **a**nti-ACTA2 (Sigma-Aldrich #C6198), anti-GPNMB (Cell Signaling Technologies #38313), anti-PODXL (R&D Systems, #AF1658), anti-CDH1 (Abcam # AB40772), anti-human Lamin A/C (Abcam, #ab108595), anti-PLVAP (Univ. of Iowa, MECA32), anti-phospho S6 (CST #2211), anti-Ki67 (ThermoFisher Scientific, # 701198), anti-human cleaved Caspase 3 (Abcam #ab2302). Biotinylated lotus tetranoglobus lectin was purchased from Fisher Scientific (#NC0370187) and visualized with streptavidin-Cy5 (ThermoFisher Scientific, #SA1011). Images were taken using a Nikon C1 confocal microscope.

### Fabrication of rapamycin nanoparticles

Rapamycin was purchased from LC Laboratories (# R5000), Poly(ethylene glycol)-block-poly(ε−caprolactone) methyl ether (PEG-PCL, PCL average Mn ∼5,000, PEG average Mn ∼5,000) and Tetrahydrofuran (THF, ≥ 99.9%) were acquired from Sigma-Aldrich. Bodipy (FL and 650/665-X, SE) were obtained from Invitrogen. A solvent evaporation method under ultrasonication was used for the encapsulation of rapamycin with PEG-PCL micelles. A 20:1 theoretical loading ratio was used for the system fabrication (10 mg of PEG-PCL diblock copolymer and 0.5 mg of rapamycin). Rapamycin and PEG-PCL were dissolved in tetrahydrofuran and added drop by drop into MillQ water under ultrasonication (Sonic Dismembrator Model 500, Fisher Scientific, Waltham, MA) at 20% amplification. The solvent was left to evaporate overnight, after which loaded micelles nanoparticles were filtered through a 0.2 um syringe filter to remove non-encapsulated aggregates in the solution. The solution was then centrifuged for 20 minutes at a rotational speed of 3500 rpm at room temperature. The preparation was stored at 4°C to avoid degradation. Nanoparticle morphology and size was analyzed using a JEOL 2010 Advanced High-Performance Transmission Electron Microscope (JEOL, MIT MRSEC, MA). The hydrodynamic radius and Zeta potential for the fabricated rapamycin-loaded nanoparticles were determined by photon correlation spectroscopy using a Zetasizer Nano ZS (Malvern Instruments, Worcester-shire, U.K.). The hydrodynamic radius was measured using a polystyrene cuvette, and the Zeta potential using a folded capillary Zeta cell. Hydrodynamic radius is reported in nm and Zeta potential in mV. Chemical analysis of functional groups of the nanoparticles was done using an FTIR spectrophotometer. Measurements for the infrared spectra of the nanoparticles were studied comparing the spectra of the reactants before and after the fabrication to observe the change in chemical composition.

### Organoid transplantation and nanoparticle delivery

Ten-week-old male NIH-Foxn1^rnu^ immunodeficient nude rats were purchased from Charles River Laboratories and maintained in the animal unit located at the Harvard Institutes of Medicine. For the surgery, the animals were anesthetized using isoflurane and weighed. The kidney was reached through a 2-cm retroperitoneal incision, and two 18-day organoids embedded in LDEV-Free hESC-qualified Matrigel were placed in the subcapsular space through a 2-mm incision in the kidney capsule. After the dorsal incision was sutured, the animals received 1ml saline and allowed to recover. Two weeks after organoid transplantation, 2μl of rapamycin-loaded nanoparticle suspensions were mixed with 10μl Matrigel and injected under the kidney capsule, approximately 2-mm away fom each organoid graft. All experiments were performed under protocols approved by the Institutional Animal Care and Use Committee at Brigham and Women’s Hospital.

### Flow cytometry

In each experiment, five organoids of each genotype were incubated with TrypLE Select (Thermo Fisher Scientific #12563011) in a shaker at 37°C for 10 min. The digestions were passed through a 40-µm cell strainer (Millipore Sigma #CLS431750), and washed with PBS. Resulting single cell suspensions were collected by centrifugation at 400*g* for 5 min. Intracellular stainings with anti-PMEL antibody HMB-45-FITC (Novus Biologicals #NBP2-34638F) and with anti-ACTA2 (Sigma-Aldrich #C6198) were performed using the PerFix-nc Kit (Beckman Coulter #B31167), for 30 min at 4 °C in supplemented PBS containing 2 mM EDTA and 2% FBS. Analysis was performed on LSRII (Becton Dickinson) equipped with three lasers. Data were collected using FacsDIVA software. Sorting gates were defined on the basis of fluorescence-minus-one control stains.

### EdU TUNEL imaging

TUNEL imaging was performed on frozen fixed graft tissue sections using the Click-iT TUNEL Alexa Fluor Imaging Assay (Invitrogen), following the manufacturer’s instructions. Briefly, the sections were incubated with anti-PMEL antibody for 1 h at room temperature, then washed and fixed with 4% PFA for 15 min at room temperature, followed by permeabilization with 0.25% Triton X-100 in PBS for 20 min at room temperature. The sections were then incubated for 1 h at 37 °C with labeling reaction mix containing TdT and nucleotide mix including EdUTP. The sections were washed and incubated in Click reaction mix including the Alexa Fluor 488 azide. After washing, the slides were incubated with Alexa 546 anti-mouse secondary antibody and stained with DAPI.

### Whole organoid transcriptome RNA sequencing and bioinformatics analysis

For each sample, total RNA was extracted from 5 organoids using the RNeasy Plus MiniKit (Qiagen). Purity was determined byA260–A280. Copy DNA (cDNA) library preparation and sequencing was performed at the Molecular Biology Core Facilities at Dan-Farber Cancer Institute. Samples were sequenced using an Illumina MiSeq sequencer and paired-end 100-bp reads generating 20-30M reads *per* sample library. The hg19 reference sequence was used for sample alignment. Hallmark, KEGG and Gene Ontology (GO) analyses for organoid comparisons were performed using Gene Set Enrichment Analysis tool (GSEA; https://www.gsea-msigdb.org) using a phenotype permutation of 1000. The false discovery rate (FDR) used to define overrepresented gene sets was q<0.25. RNA-sequencing of 18 fresh/frozen kidney angiomyolipomas and 4 normal kidneys and expression data from this set were combined to publicly available data from 10 additional kidney angiomyolipomas and 4 normal kidneys (42) for downstream analyses. The RNA-sequencing method was previously described (26) following the manufacturer’s instructions.

### Cytochrome C ELISA

Blood samples were collected from the tail vain of RNU rats prior to the transplantation procedure and on days 3 and 7 after surgery. Cytochrome C was detected the serum samples using the Human Cytochrome C ELISA Kit (ab221832) following the manufacturer’s instructions.

### Statistics

Student’s *t*-test, one-way ANOVA and two-way ANOVA were used. Where applicable, normal distribution of the data was determined using D’Agostino–Pearson and Shapiro– Wilk normality tests. Statistical analysis was performed using Prism 7 for Mac OS (GraphPad Software, Inc.).

## Data Availability

Source data are provided with this paper. The minimal datasets necessary to interpret, replicate and build upon the methods or findings reported in this article are fully available in the online version. All of the data are available from the corresponding author upon reasonable request.

## Acknowledgements

This study was supported by grant funding from the National Institutes of Health (NIH) (R21AG058159 to DRL, R01DK124301 to DRL and R01NS113591 to MS) and the Intellectual and Developmental Disabilities Center at Boston Children’s Hospital (BCH IDDRC; U54HD090255).

