## Supplementary Data for "A Tissue-Bioengineering Strategy for Modeling Rare Human Kidney Diseases *In Vivo*"

Number of DEGs and transcription factors identified in the comparisons *TSC2*$^{-/-}$ *vs. TSC2*$^{+/+}$ renal organoids and *TSC2*$^{-/-}$ *vs. TSC2*$^{-/+}$ renal organoids

| **Differential gene expression DESeq2 analysis** | ***TSC2 -/-* vs *TSC2 +/+* kidney organoids** | | ***TSC2 -/-* vs *TSC2 +/-* kidney organoids** | |
|---|---|---|---|---|
| **All DEGs (FDR<0.05 and $log_2$>2/$log_2$<-2)** | **477 ($log_2$ >2)** | **362 ($log_2$<-2)** | **480 ($log_2$ >2)** | **363 ($log_2$<-2)** |
| **All TFs FDR<0.05 and $log_2$>2/$log_2$<-2)** | **59 ($log_2$ >2)** | **30 ($log_2$<-2)** | **60 ($log_2$ >2)** | **31 ($log_2$<-2)** |

**Supplementary Table 1.** Pairwise differential gene expression analyses in *TSC2*$^{-/-}$ kidney organoids compared to *TSC2*$^{+/+}$ and *TSC2*$^{+/-}$ kidney organoids. Numbers of all upregulated/downregulated genes and transcription factors for adjusted p-value/FDR<0.05 and $log_2$ fold >2/$log_2$ fold <-2 for expression FPKM values are presented.

**Supplementary figure 1. a** Differentiation schematic and representative immunofluorescence images showing the effect of a 24-h retinoic acid pulse on Day 13 on formation of glomeruli containing $PODXL^{+}$ cells. **b** Quantification of the number of glomeruli *per* field recorded with the microscope (20X). Bar graph shows mean ± SD, n = 4 experiments of three wells each. **P*<0.05. Scale bar 50μm.

**Supplementary figure 2. a** Morphological appearance of AML cells in $TSC2^{-/-}$ renal organoids compared to $TSC2^{+/+}$ renal organoid and kidney fibroblasts. **b** Scattered distribution of AML cells compared to the peritubular organization of fibroblasts in normal and fibrotic $TSC2^{+/+}$ renal organoids. Fibrosis was induced by incubation with IL1β for 96h. Scale bars 25μm.

**Supplementary figure 3. a** Expression of housekeeping genes in RNAseq samples of $TSC2^{+/+}$, $TSC2^{-/+}$, and $TSC2^{-/-}$ renal organoids, compared to kidney angiomyolipomas and normal kidneys. **b**, **c** Tables showing Hallmark gene pathways identified by GSEA analysis of $TSC2^{+/+}$ *vs.* $TSC2^{-/-}$ (**b**) and $TSC2^{-/+}$ *vs.* $TSC2^{-/-}$ (**c**) renal organoid gene expression comparisons.

**Supplementary figure 4. a** Flow cytometry gating strategy used to identify $ACTA2^{+}$ $PMEL^{+}$ cells in $TSC2^{-/-}$ renal organoids. **b**, **c** Single color controls lacking anti-ACTA2 antibody (**b**), and anti-PMEL antibody (**c**). The numbers indicate representative cell frequencies.

**Supplementary figure 5. a** Representative brightfield images showing *TSC2*$^{-/-}$ renal organoid cyst growth. Scale bars 500μm. **b** Representative whole mount immunofluorescence showing the presence of cysts lined by CDH-expressing TECs in *TSC2*$^{-/-}$ renal organoids.

**Supplementary figure 6. a** Schematic representation of the process of Rapa-nanoparticle fabrication. **b** Visualization of nanoparticle morphology using transmission electron microscopy. **c** TEM micrograph analysis showing the symmetric spherical shape and narrow size distribution of 12 nm with no sign of bulk participations for the Rapamycin-loaded PEG-PCL nanoparticles. **d** Represenative immunofluorescence microscopy images showing Ki67 in primary mouse TECs incubated with control DMEM medium + 5% FBS (Control), DMEM medium + empty nps (Empty-Nps), DMEM medium + rapamycin alone (Rapamycin), or DMEM medium + Rapa-Nps (Rapa-Nps). **d** Quantification of Ki67 expression, bars graph represents mean ± SD, n = 3 independent experiments. ***P*<0.01, ****P*<0.005. Representative fluorescence imaging showing the viability of Day-21 *TSC2*$^{-/-}$ renal organoids exposed to rapamycin nanoparticles, at 0h and 48h of incubation. The Live cell staining is shown in the green channel, while the Dead cell staining is shown in the red channel. **b** Quantification of Live cell and Dead cell signal at both time points studied. Four wells containing one organoid each were measured. Floating bars graph represents mean ± SD, n = 4 organoids. **P*<0.05. Scale bars, 500µm.

### Supplementary Figure 1

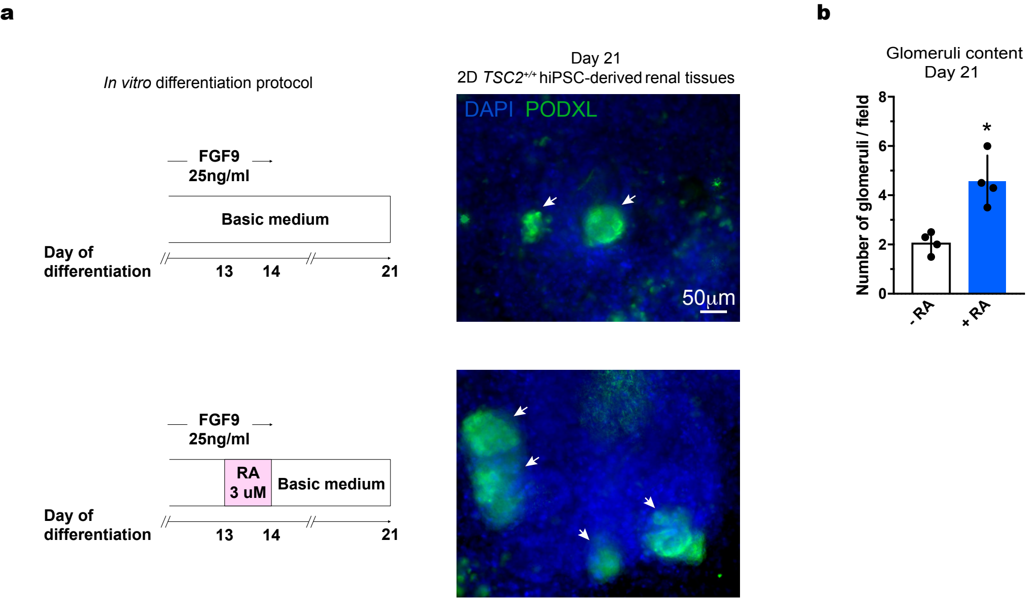

### Supplementary Figure 2

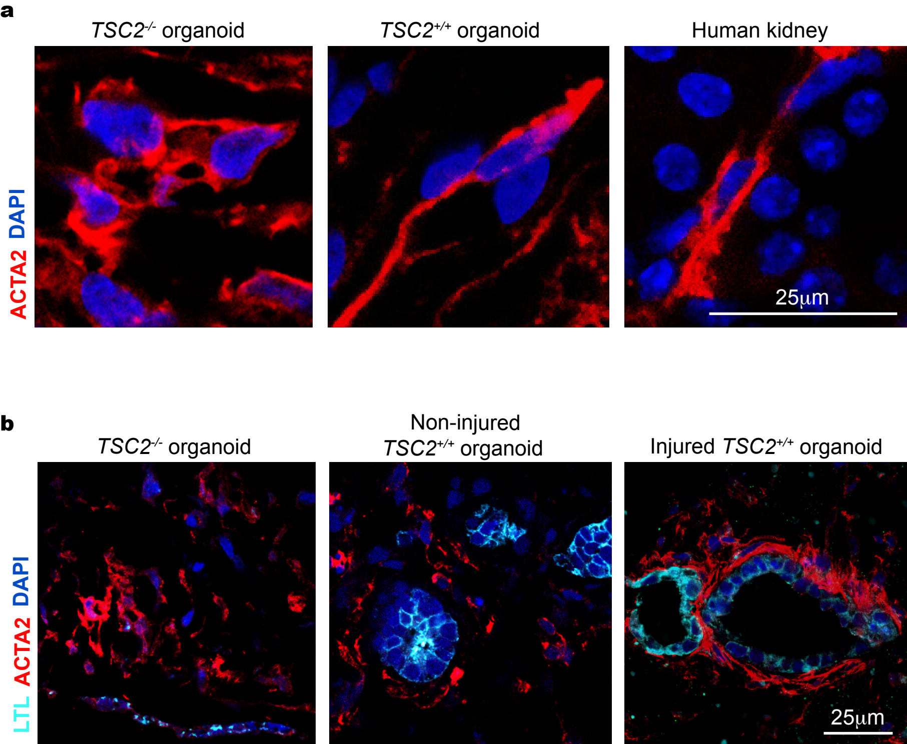

### Supplementary Figure 3

**a**

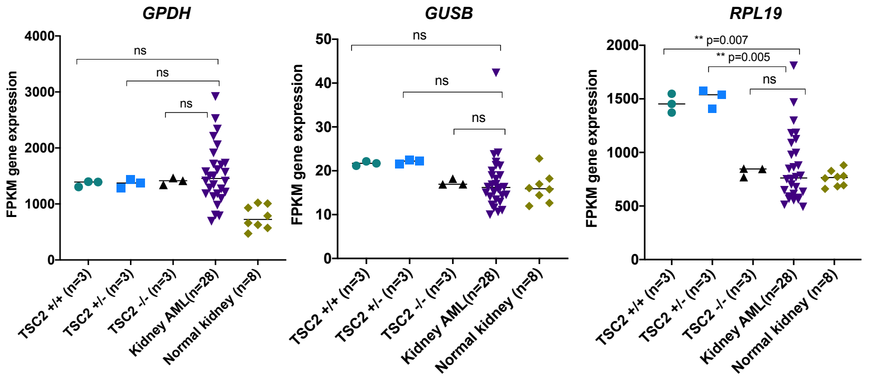


**b**

| NAME | ES | NES | NOM p-val | FDR q-val | FWER p-val |
|---|---|---|---|---|---|
| HALLMARK_REACTIVE_OXYGEN_SPECIES_PATHWAY | 0.39871022 | 1.7069229 | 0 | 0.05499997 | 0 |
| HALLMARK_COMPLEMENT | 0.59225315 | 1.5513009 | 0 | 0.10523207 | 0.137 |
| HALLMARK_ANGIOGENESIS | 0.58937246 | 1.4004242 | 0 | 0.14682235 | 0.363 |
| HALLMARK_XENOBIOTIC_METABOLISM | 0.4329361 | 1.3941399 | 0 | 0.15006849 | 0.363 |
| HALLMARK_KRAS_SIGNALING_UP | 0.47650862 | 1.4073737 | 0 | 0.150379 | 0.363 |
| HALLMARK_INFLAMMATORY_RESPONSE | 0.4672174 | 1.3784195 | 0 | 0.15125427 | 0.465 |
| HALLMARK_ALLOGRAFT_REJECTION | 0.5906101 | 1.466401 | 0 | 0.1587153 | 0.196 |
| HALLMARK_ESTROGEN_RESPONSE_LATE | 0.3445101 | 1.3827776 | 0 | 0.16000482 | 0.465 |
| HALLMARK_INTERFERON_GAMMA_RESPONSE | 0.49505466 | 1.3540543 | 0 | 0.1606237 | 0.465 |
| HALLMARK_PEROXISOME | 0.2900785 | 1.4142873 | 0 | 0.16400446 | 0.363 |
| HALLMARK_TNFA_SIGNALING_VIA_NFKB | 0.3182069 | 1.3261827 | 0 | 0.1682529 | 0.51 |
| HALLMARK_INTERFERON_ALPHA_RESPONSE | 0.48019886 | 1.3278441 | 0 | 0.17634226 | 0.51 |
| HALLMARK_IL2_STAT5_SIGNALING | 0.35907206 | 1.3062559 | 0 | 0.18003435 | 0.602 |
| HALLMARK_APICAL_JUNCTION | 0.36009854 | 1.3188215 | 0 | 0.18120329 | 0.602 |
| HALLMARK_COAGULATION | 0.64518267 | 1.424169 | 0 | 0.18217193 | 0.363 |
| HALLMARK_FATTY_ACID_METABOLISM | 0.38412678 | 1.2796404 | 0 | 0.1853368 | 0.646 |
| HALLMARK_IL6_JAK_STAT3_SIGNALING | 0.6289876 | 1.4391141 | 0 | 0.18589109 | 0.251 |
| HALLMARK_BILE_ACID_METABOLISM | 0.37456596 | 1.4672211 | 0 | 0.19328701 | 0.196 |
| HALLMARK_APOPTOSIS | 0.27065846 | 1.226526 | 0 | 0.20427474 | 0.742 |
| HALLMARK_ADIPOGENESIS | 0.26161525 | 1.2311289 | 0.10412574 | 0.20908016 | 0.742 |
| HALLMARK_UV_RESPONSE_DN | 0.28318453 | 1.239677 | 0.22857143 | 0.21439168 | 0.742 |
| HALLMARK_EPITHELIAL_MESENCHYMAL_TRANSITION | 0.41336355 | 1.1974243 | 0 | 0.21484017 | 0.786 |

**c**

| NAME | ES | NES | NOM p-val | FDR q-val | FWER p-val |
|---|---|---|---|---|---|
| HALLMARK_REACTIVE_OXYGEN_SPECIES_PATHWAY | 0.39871022 | 1.6877129 | 0 | 0.05399998 | 0 |
| HALLMARK_COMPLEMENT | 0.59225315 | 1.5637503 | 0 | 0.12149391 | 0.168 |
| HALLMARK_ANGIOGENESIS | 0.58937246 | 1.4073582 | 0 | 0.13223304 | 0.305 |
| HALLMARK_XENOBIOTIC_METABOLISM | 0.4329361 | 1.4113219 | 0 | 0.13340722 | 0.305 |
| HALLMARK_PEROXISOME | 0.2900785 | 1.4120195 | 0 | 0.14333321 | 0.305 |
| HALLMARK_INTERFERON_GAMMA_RESPONSE | 0.49505466 | 1.3601393 | 0 | 0.14820744 | 0.459 |
| HALLMARK_ESTROGEN_RESPONSE_LATE | 0.3445101 | 1.3781137 | 0 | 0.14829785 | 0.459 |
| HALLMARK_INFLAMMATORY_RESPONSE | 0.4672174 | 1.3823795 | 0 | 0.14841431 | 0.459 |
| HALLMARK_COAGULATION | 0.64518267 | 1.4263659 | 0 | 0.15324047 | 0.257 |
| HALLMARK_KRAS_SIGNALING_UP | 0.47650862 | 1.4144634 | 0 | 0.15609506 | 0.305 |
| HALLMARK_ALLOGRAFT_REJECTION | 0.5906101 | 1.4698694 | 0 | 0.15863593 | 0.214 |
| HALLMARK_TNFA_SIGNALING_VIA_NFKB | 0.3182069 | 1.3359405 | 0 | 0.16024484 | 0.504 |
| HALLMARK_IL2_STAT5_SIGNALING | 0.35907206 | 1.3176168 | 0 | 0.16463935 | 0.504 |
| HALLMARK_IL6_JAK_STAT3_SIGNALING | 0.6289876 | 1.444351 | 0 | 0.16477522 | 0.214 |
| HALLMARK_INTERFERON_ALPHA_RESPONSE | 0.48019886 | 1.3389647 | 0 | 0.1678337 | 0.504 |
| HALLMARK_APICAL_JUNCTION | 0.36009854 | 1.3079895 | 0 | 0.17421544 | 0.558 |
| HALLMARK_FATTY_ACID_METABOLISM | 0.38412678 | 1.2886324 | 0 | 0.17870884 | 0.66 |
| HALLMARK_UV_RESPONSE_DN | 0.28318453 | 1.269903 | 0.19842829 | 0.17953376 | 0.66 |
| HALLMARK_BILE_ACID_METABOLISM | 0.37456596 | 1.4786377 | 0 | 0.19351456 | 0.214 |
| HALLMARK_ADIPOGENESIS | 0.26161525 | 1.2390532 | 0.10136452 | 0.19938481 | 0.709 |
| HALLMARK_EPITHELIAL_MESENCHYMAL_TRANSITION | 0.41336355 | 1.2075171 | 0 | 0.20924948 | 0.759 |
| HALLMARK_APOPTOSIS | 0.27065846 | 1.2129807 | 0 | 0.21453014 | 0.759 |

### Supplementary Figure 4

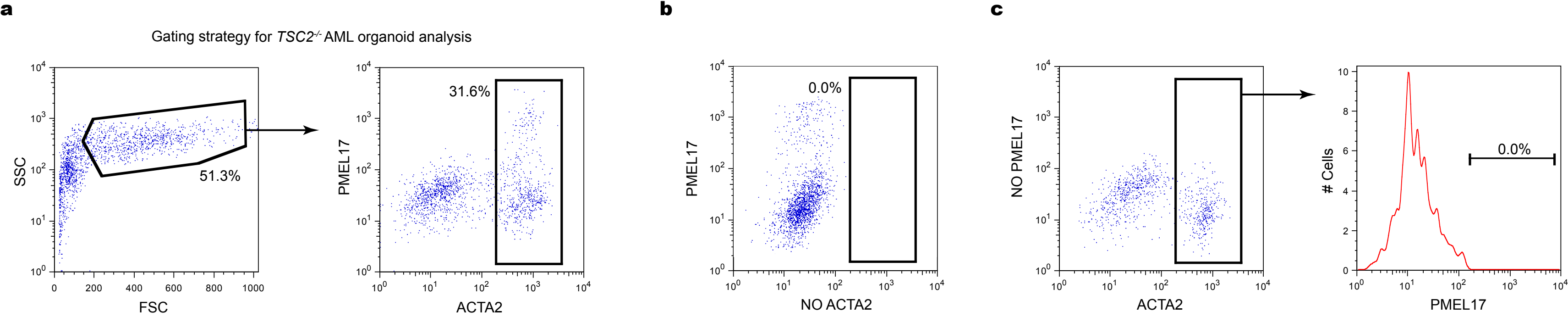

### Supplementary Figure 5

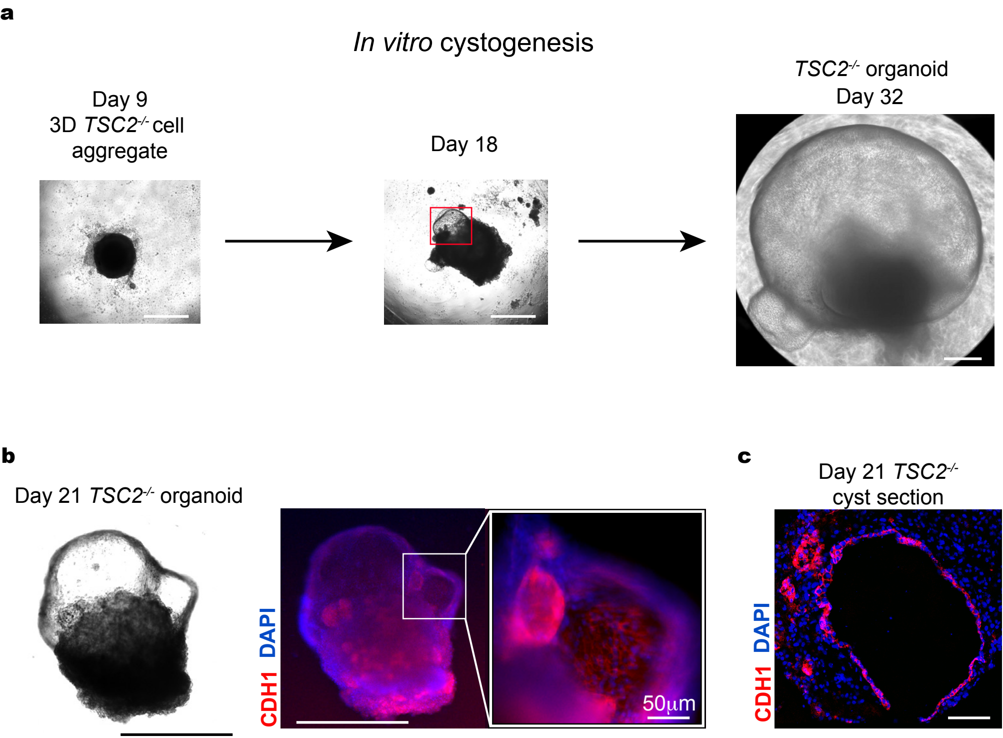

### Supplementary Figure 6

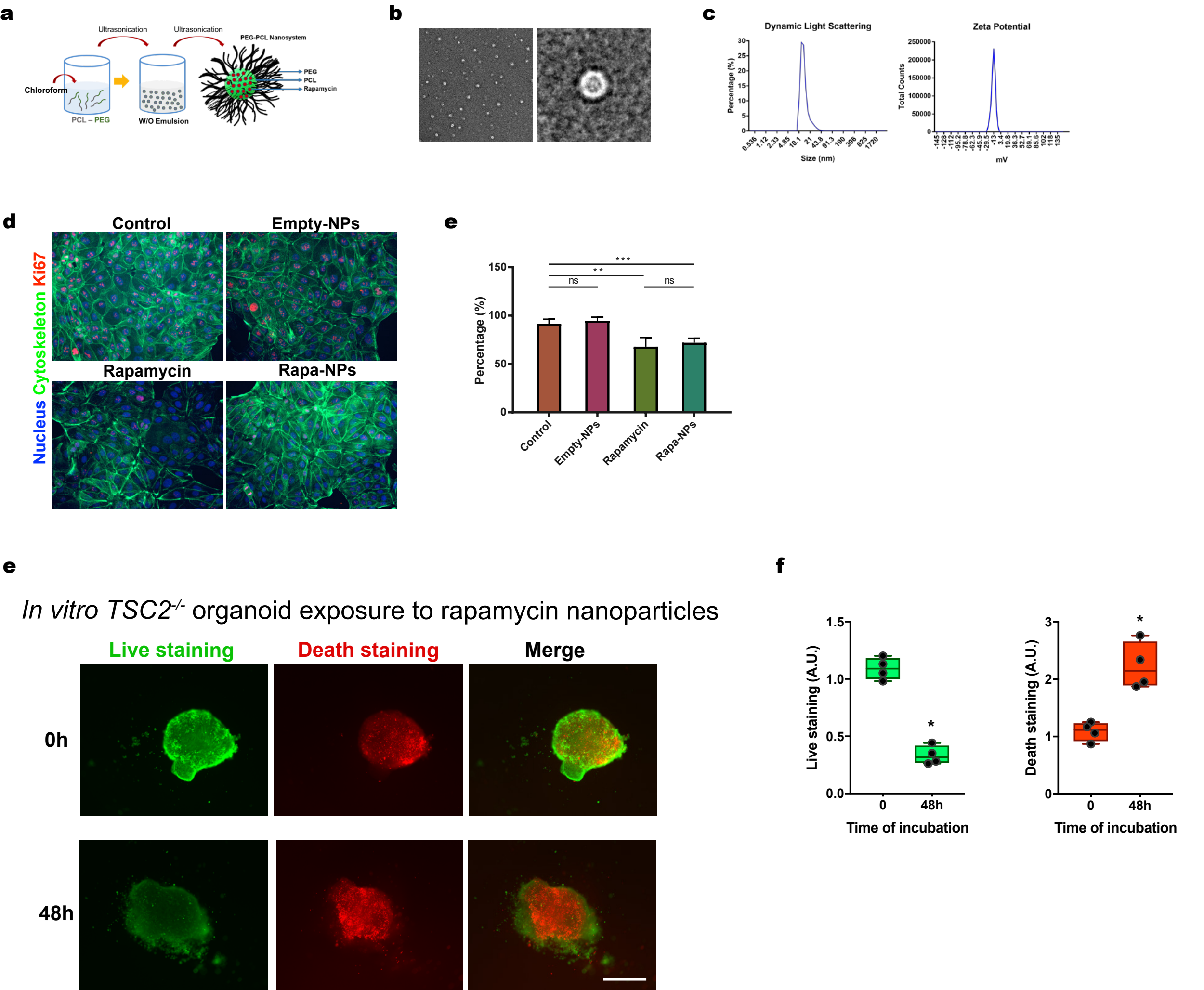

**Supplementary Table.** List of Taqman RT-PCR primers used.

| Taqman Assay ID | Human Gene | Catalog # |
|---|---|---|
| Hs00232731_m1 | *SIX2* | 4331182 |
| Hs04285637_m1 | *SALL1* | 4331182 |
| Hs01015256_g1 | *PAX8* | 4351372 |
| Hs00232144_m1 | *LHX1* | 4331182 |
| Hs01001602_m1 | *HNF1B* | 4331182 |
| Hs01103751_m1 | *WT1* | 4331182 |
| Hs03023943_g1 | *ACTB* | 4331182 |
